# Connexin43 controls N-cadherin transcription during collective cell migration

**DOI:** 10.1101/114371

**Authors:** Maria Kotini, Elias H. Barriga, Jonathan Leslie, Marc Gentzel, Alexandra Schambony, Roberto Mayor

## Abstract

Connexins are the primary components of gap junctions, providing direct links between cells in many physiological processes, including cell migration and cancer metastasis. Exactly how cell migration is controlled by gap junctions remains a mystery. To shed light on this, we investigated the role of Connexin43 in collective cell migration during embryo development using the neural crest, an embryonic cell population whose migratory behavior has been likened to cancer invasion. We discovered that Connexin43 is required for contact inhibition of locomotion by directly regulating the transcription of N-cadherin. For this function, the Connexin43 carboxy tail interacts with Basic Transcription Factor 3, which mediates its translocation to the nucleus. Together, they bind to the n-cad promotor regulating n-cad transcription. Thus, we uncover an unexpected, gap junction-independent role for Connexin43 in collective migration that illustrates the possibility that connexins, in general, may be important for a wide variety of cellular processes that we are only beginning to understand.

**Highlights:**

- Cx43 regulates collective directional migration of neural crest cells
- Cx43 carboxy tail controls cell polarity via n-cad regulation
- Cx43 carboxy tail localises at the nucleus and that depends on BTF3
- BTF3 and Cx43 carboxy tail directly interact to bind and regulate n-cad promoter activity

## Introduction

Connexins, the protein components of gap junctions (GJs), have been established as pivotal molecules in morphogenesis (Liu et al., 2012; Mathias et al., 2010) and cell migration during embryonic development (Cina et al., 2009; Elias et al., 2007; Francis et al., 2011) and cancer metastasis (Ogawa et al., 2012; Stoletov et al., 2013). Yet the way that connexins regulate these processes is not fully understood. While the activity of connexins as channels has been shown critical for the regulation of tissue homeostasis (Alexander and Goldberg, 2003; Plum et al., 2000), it is not required for cell migration (Cina et al., 2009; Elias et al., 2007), an important aspect that has been emphasised in various cell types for Cx43, the most abundant and conserved connexin among vertebrates, (Elias et al., 2007, 2010; Homkajorn et al., 2010). Attemps to build a complete picture of how Cx43 functions in cell migration have been complicated by the fact that Cx43 possesses several functions. Indeed, Cx43 can work as a channel (Niessen et al., 2006) or a hemi-channel (Bruzzone et al., 2001; Ebihara et al., 1999), as an adhesive molecule (Elias et al., 2007, 2010; Elzarrad et al., 2008) and can also be involved in intracellular signaling (Cina et al., 2009). Additionally, a number of molecules interact with Cx43 (Giepmans et al., 2001; Saidi Brikci-Nigassa et al., 2012) and have been implicated in the regulation of the Cx43 activities (Ai et al., 2000; Kwak et al., 1995; Solan and Lampe, 2005). Furthermore, it has been shown that Cx43 generates small carboxy terminal isoforms (Salat-Canela et al., 2014; Smyth and Shaw, 2013), yet the biological function of these isoforms remains unknown.

Numerous studies support the notion that Cx43 is involved in cell migration via different activities (Liu et al., 2012; Mendoza-Naranjo et al., 2012; Waldo et al., 1999; Zhang et al., 2014). Interestingly, one aspect that is common to a number of cell types, including neurons, astrocytes, cardiac neural crest and metastatic cancer cells, is that loss of function of Cx43 leads to an effect on cell protrusions such as lamellipodia or filopodia and a concurrent loss in cell polarity (Elias et al., 2007; Homkajorn et al., 2010; Xu et al., 2006; Stoletov et al., 2013). In certain instances, where directional motion is involved, this loss of cell polarity may contribute to the defective migration (Elias et al., 2010; Francis et al., 2011; Xu et al., 2006). Despite this, exactly how Cx43 controls cell migration, polarity and protrusions remains unknown.

Collective cell migration is dependent on cell-cell cooperation (Friedl and Gilmour, 2009; Mayor and Ettienne-Manvile, 2016), but our knowledge on the molecular mechanism of this cell cooperation remains incomplete. For example, we know that adherens junctions and specifically N-cadherin (N-cad) is critical for establishing cohesiveness of the cells and cell polarity in a collective (Theveneau et al., 2010; Kuriyama et al., 2014). Interestingly, it has been shown that Cx43 can modify the levels of N-cad at the membrane (Giepmans, 2006; Xu et al., 2001), but the mechanism driving this regulation remains unknown.

To disentangle this problem we used *Xenopus* cranial neural crest (NC), which provides a well-established model for directional motion of cell collectives (Theveneau and Mayor, 2013). An essential phenomenon required for NC migration is contact inhibition of locomotion (CIL), the process by which cells polarise away from cell contacts within a cell collective (Carmona-Fontaine et al., 2008; Stramer et al., 2013). This contact dependent polarity is essential for collective motion and requires the formation of a transient cell-cell adhesion junction, with N-cad being a crucial element, which needs to be tightly regulated as excess or deficiency of N-cad is sufficient to impair NC migration (Scarpa et al., 2015; Theveneau et al., 2010). However, exactly how the levels of n-cad are controlled during collective behavior is poorly understood.

Here we propose a link between cell polarity and the regulation of N-cad levels by Cx43. We show that Cx43, a molecule primarily known for its membrane-linked activities, uses its tail isoform to control morphogenetic movements via transcriptional regulation of n-cad, an activity which is independent from its function as channel in the cell membrane. Moreover, we identify its mechanism of action by showing that Cx43 regulation of n-cad is due to a direct interaction with the basic transcription factor 3 (BTF3) and polymerase II. Adittionally we show that this complex directly binds to the n-cad promoter to modulate its transcription. Furthermore, we show that this novel activity of Cx43 as a regulator of n-cad is conserved between amphibian and mammalian cells.

## Results

### Cx43 modulates NC migration *in vivo* and *in vitro* via the regulation of cell polarity

Previous studies have shown that Cx43 is expressed in mouse cardiac NC and that aberrant Cx43 expression leads to migration deficiency (Ewart et al., 1997; Waldo et al., 1999). In order to study the cellular and molecular roles of Cx43 in migration we used the cephalic NC from *Xenopus laevis* embryos because it is an amenable system for *in vivo* and *in vitro* characterisation of cell migration. In *Xenopus,* cephalic NC cells can be identified by the expression of the transcription factor *twist* (Linker et al., 2000). Using antibody staining, we found that Cx43 is expressed in the cephalic NC migratory streams as compared with the NC marker *twist* (Figure 1A). In addition, Cx43 is also expressed at the neural tube and the presumptive eye (Figure 1A). In other systems, Cx43 is present in two forms: the full length molecule (Cx43FL), which is a component of GJs, and the smaller carboxy terminal fragments (Cx43iso) that are not part of the GJ structures (Salat-Canela et al., 2014; Smyth and Shaw, 2013). To identify whether these isoforms are present in *Xenopus,* we performed western blot analysis of Cx43 at different developmental stages. Our analysis revealed that Cx43FL (between 35kDa and 55 kDa) is expressed throughout all the stages analysed (Figure 1B, stages 16 to 25), while the expression of Cx43iso (between 15kDa and 25kDa) increases just before the NC start its migration (Figure 1B, stage 18 onwards). These results were confirmed by analysing Cx43 mRNA using RT-PCR, which shows that Cx43 mRNA is present in dissected NC and in whole embryos at stage 18 (Figure S1A).

**Figure 1.**
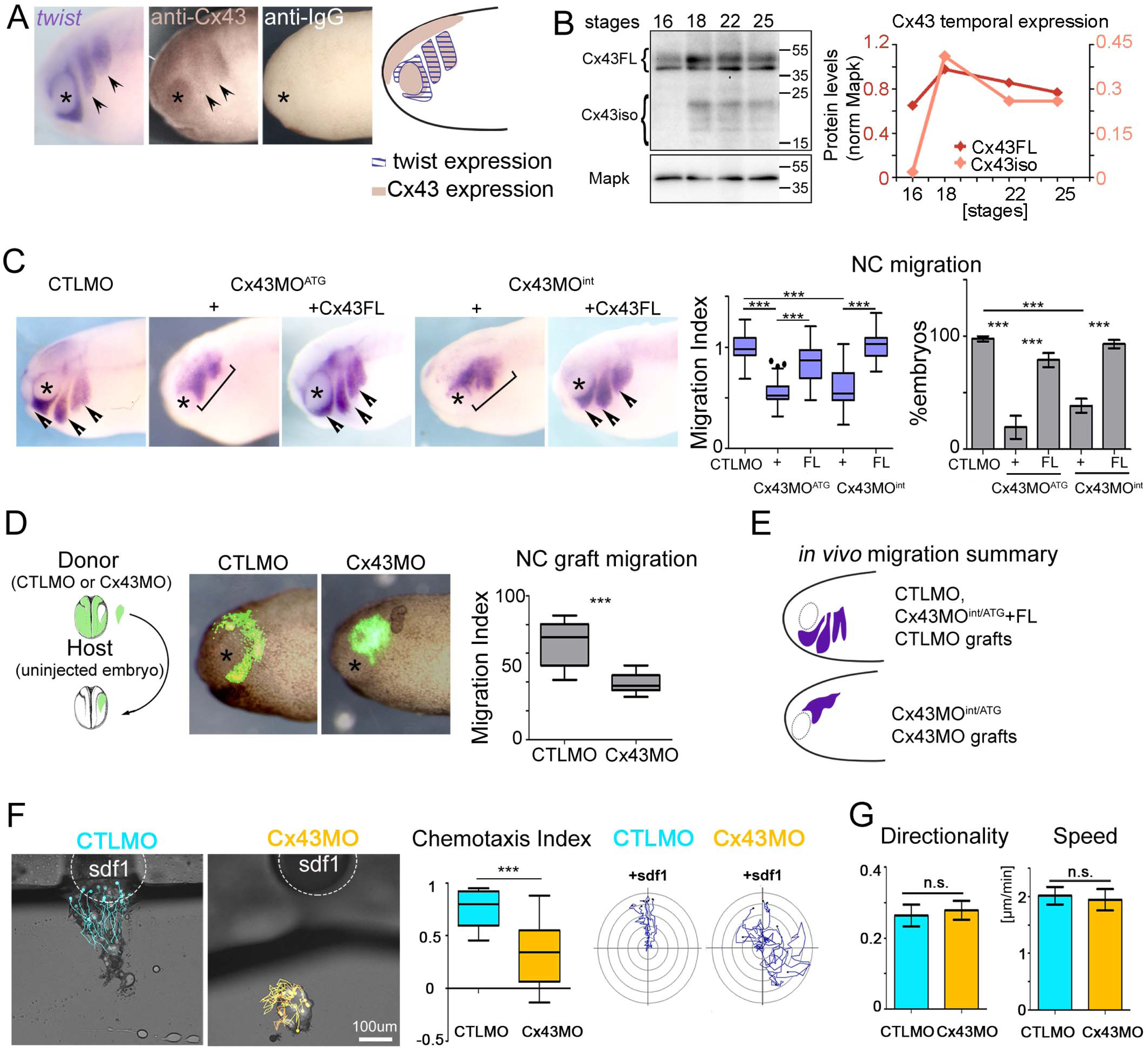
Cx43 controls NC migration. (A) *twist* ISH and Cx43 immunostaining for st24 embryo; IgG control antibody; arrows indicate NC, asterisk marks the eye. Drawing: Cx43 and twist expression summary. (B) WB for Cx43 and Mapk. Cx43 temporal expression at neurula stages. (C) *twist* ISH for st24 embryos, arrows show normal and brackets impaired migration. Quantification of migration index (n_CTLMO_=31, n_Cx43MO_^ATG^=37, n_Cx43MO_^ATG^+_Cx43FL_=30, n_Cx43MO_^int^=33, n_Cx43MO_^int^_+Cx43FL_=19, N = 3, median ± interquartile range, p***<0.001) and % of embryos (N=3, mean ± SE, p***<0.001). (D) Transplant assay summary and experiment. Migration index of grafted NC (n=15, N=3, median ± interquartile range, p***<0.001). (E) *In vivo* migration summary. (F) Cell tracks of NC explants towards SDF-1. Tracks for single explants and their chemotaxis index (median ± quartile, n=15, N=3, p***<0.001). (G) Directionality and velocity of NCs (n_CTLMO_=36, n_Cx43MO_=31, N=4, mean ± SE, ns=non-significance). See also Figure S1.

To test whether Cx43 is required for NC migration, we blocked translation of Cx43 using two antisense morpholino variants, Cx43MO^ATG^and Cx43MO^int^ (see Star Methods, Figure S1C). Injection of any of these two MOs leads to a strong inhibition of NC migration when compared to the effect of a control MO (CTLMO) (Figure 1C; S1B). Moreover, co-injection of either of the MOs with human cx43FL mRNA, that does not bind the MOs, showed a rescue in NC migration confirming the specificity of these treatments (Figure 1C). To test if the observed effect is tissue-autonomous, we transplanted NC from CTLMO- or Cx43MO-injected embryos into uninjected host embryos (Figure 1D). Only embryos with Cx43MO transplanted NC showed an inhibition in migration, indicating that Cx43 is autonomously required in the NC. Importantly, our observation on migration is not an indirect result of affecting NC induction, since Cx43 downregulation does not affect the early expression of the NC marker *snail2* (Mayor et al., 1995), as shown by *in situ* hybridisation (ISH) (Figure S1D).

To assess how Cx43 influences NC migration, we analysed the effect of its loss-of-function *in vitro.* In a first step, we analysed the response of the NC population towards an external source of the chemoattractant SDF-1 (Theveneau and Mayor, 2011). In comparison to CTLMO explants, Cx43MO explants failed to migrate towards SDF-1 (Figure 1F; Supplemental Movie S1). Importantly, analysis of speed and directionality of individually migrating cells show that cell motility is not affected in Cx43MO cells (Figure 1F; Supplemental Movie S2, Figure S1E). Together these results support the notion that Cx43 is involved in the collective response required for NC migration, but not in single cell motility.

To examine how loss of Cx43 might affect cells during collective migration, we analysed the effect of Cx43MO on cell morphology. It has been previously shown that NCs form large protrusions away from cell contacts and that this polarity is required for correct NC migration (Carmona-Fontaine et al., 2008; Scarpa et al., 2015). Visualisation of actin and α-tubulin cytoskeleton showed shorter protrusions in Cx43MO cell clusters (Figure 2A, B). Consistently, analysis of cell shape demonstrates a circular and less polarised phenotype in Cx43MO cells when compared to CTLMO cells (Figure 2C). An additional way to test cell polarity is the localisation of Lamellipodin (Lpd), a regulator of Rac1 activity at the cell protrusions (Law et al., 2013). In CTLMO cells, Lpd-GFP showed higher levels at the tip of the lamella and lower levels in the cytosol and contact points, as previously described (Law et al., 2013). In contrast, in Cx43MO cells, Lpd-GFP levels were lower at the free edge and higher at the contact point, confirming that cell polarity is disrupted (Figure 2D). These observations support the involvement of Cx43 in contact-dependent polarity, and suggest an implication for contact inhibition of locomotion (CIL), which is an essential activity for directional collective cell migration (Stramer and Mayor, 2017). We tested CIL on NC cells and on NC explants. First, NC cells undergoing CIL move away from each other upon collision after a short time (10min), generating small collision angles (0^o^ corresponds to perfect repulsion). Analysis of cell collisions indicated an increase in collision duration and collision angles in Cx43 depleted NCs compared to CTLMO cells, showing an effect on CIL (Figure 2E; Supplemental Movie S3). Second, when two explants of cells that exhibit CIL are cultured a short distance apart they never overlap, as it was observed for CTLMO explants, whereas Cx43MO explants showed an increase in their area of overlap, proving an impaired CIL response (Figure 2F). Third, one of the consequences of CIL is that it promotes cell dispersion. Indeed, CTLMO cells dispersed after 4-6hrs of culture, while cells injected with Cx43MO displayed a delay in cell dispersion(Figure 2G; Supplemental Movie S4). Importantly, Cx43MO explants were able to regain their momentum and show similar levels of dispersion to CTLMO when cultured for longer times (>10hrs). Taken together, our analysis *in vivo* and *in vitro* showed that perturbed Cx43 expression inhibits migration by altering cell polarity during CIL.

**Figure 2.**
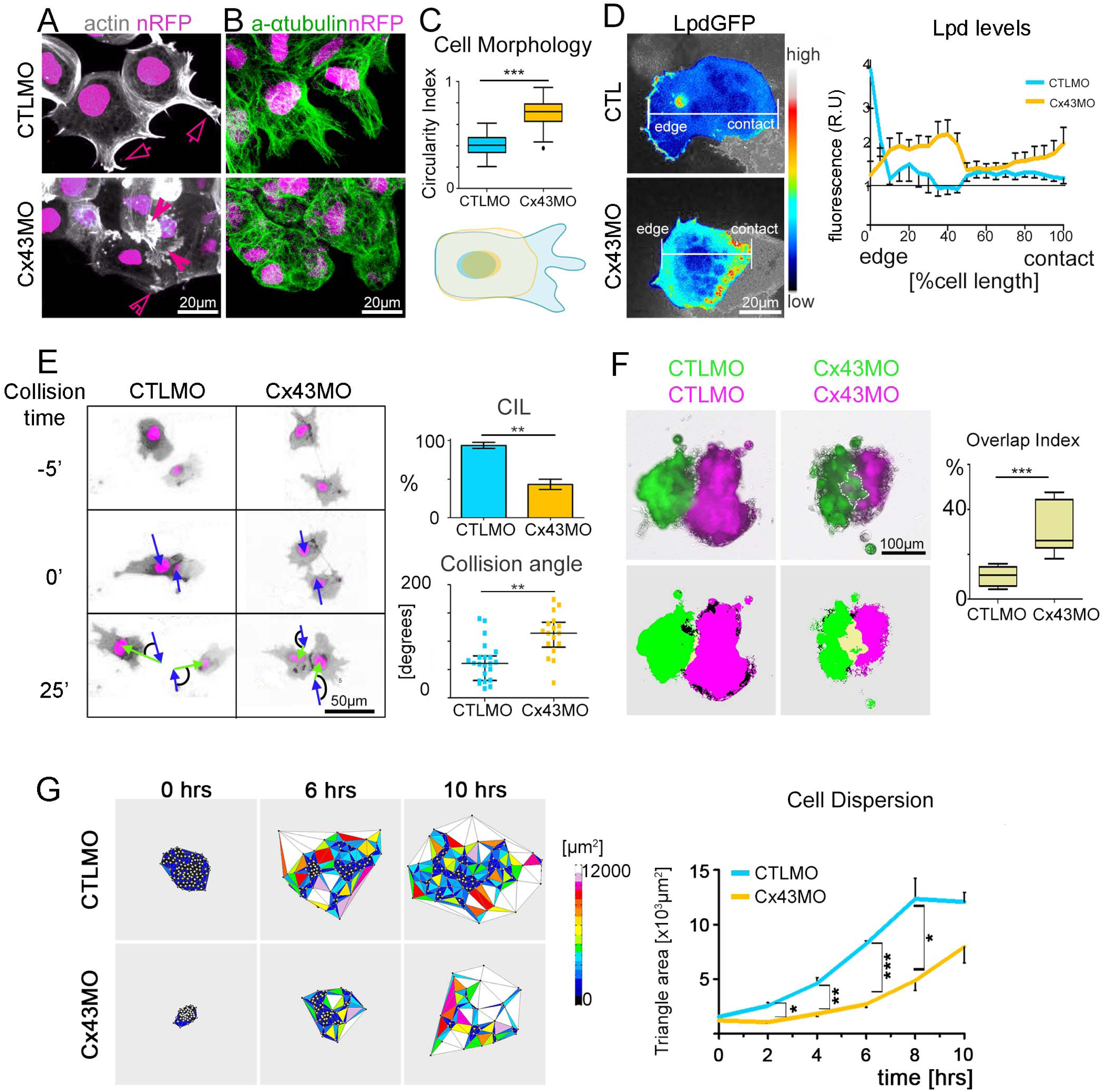
Cx43 modulates cell polarity. (A) NC cell morphology visualised by lifeactinGFP (arrows mark normal and arrowheads irregular protrusions) and by (B) Immunostaining for α-tubulin. (C) Circularity index and cell morphology diagram (n_CTLMO_=58, n_Cx43MO_=39, N=3, mean ± SE, p***<0.001). (D) Heatmap of Lpd-GFP NCs. Lpd-GFP fluorescence normalised to background levels along cell length (n=8, N=3, mean ± SE). (E) Single cell collisions (0 marks collision). Blue arrow: velocity before collision; green arrow: velocity after collision; the angle between these vectors is shown in black. Quantification of CIL (n=23, N=5, mean ± SE, p**<0.01) and of collision angle (median ± interquartile range, p**<0.01). (F) Explant confrontation assay (bottom thresholded images of top panels) and quantification of overlap index (n_CTLMO_=12, n_Cx43MO_=11, N=3 median ± interquartile range, p***<0.001). (G) Cell dispersion analysis using Delauney triangulation and its quantification (n=15, N=3, mean ± SE, p***<0.001, p**<0.01, p*<0.05).

### Cx43 regulates n-cad expression

In order to understand how Cx43 affects cell polarity we decided to investigate other molecules linked to this process. N-cad, for example, is present at cell contacts during NC migration and is critical for cell polarisation during CIL and collective migration (Theveneau et al., 2010; Kuriyama et al., 2014). Interestingly, immunostaining of NC cells revealed that in comparison to control conditions, in which N-cad is expressed and localised at cell contacts, Cx43MO explants show an N-cad reduction at cell contacts (Figure 3A). This observation was confirmed by western blot analysis that shows a decrease in the overall N-cad protein levels in Cx43MO embryo lysates when compared to CTLMO lysates (Figure 3B).

**Figure 3.**
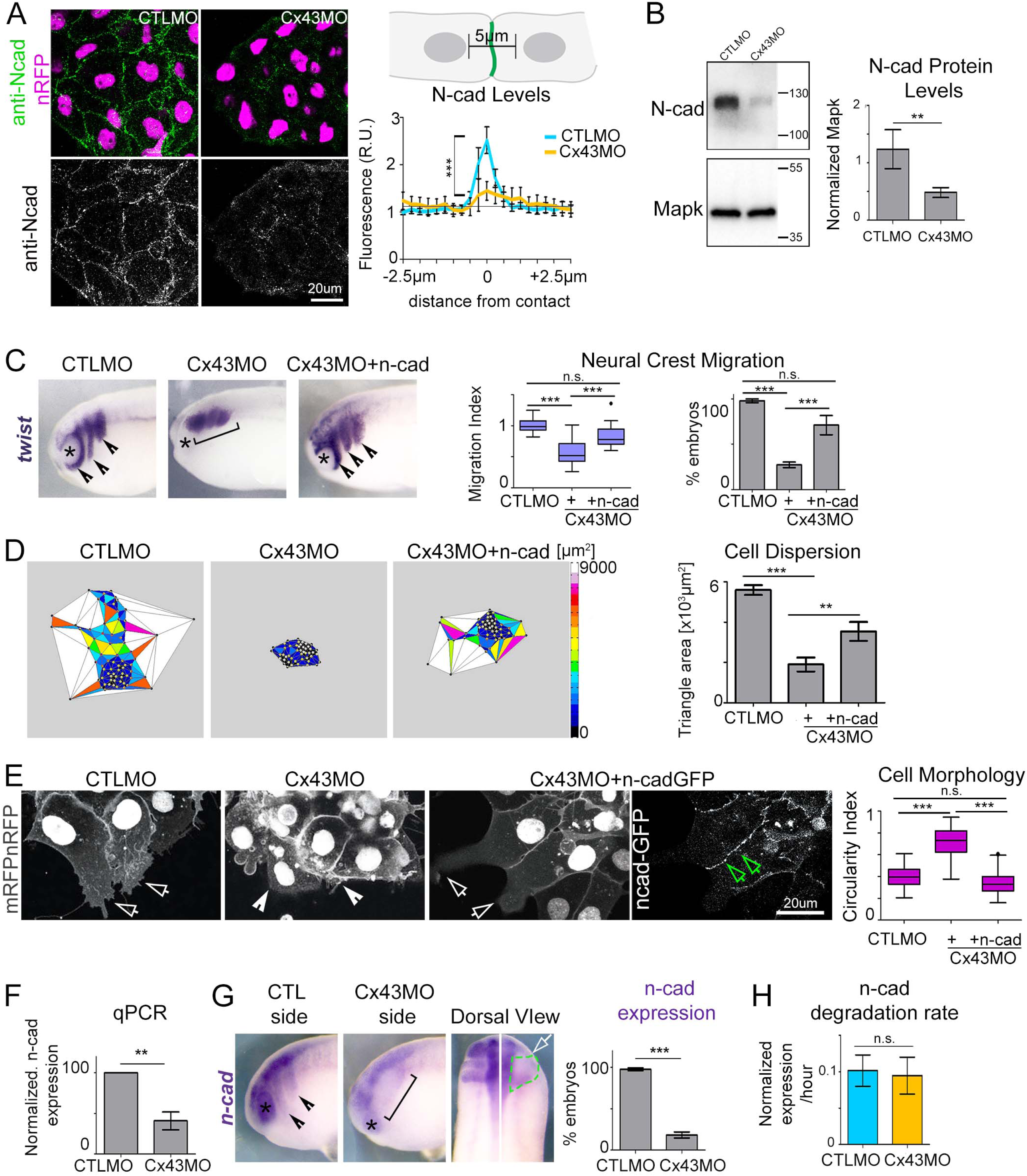
Cx43 regulates n-cad expression. (A) Immunostaining for N-cad. Summary of N-cad analysis. N-cad levels, 0 represents contact point (n=15, N=3, mean± SE, p***<0.001). (B) WB for N-cad and Mapk. N-cad levels normalised to Mapk (N=5, mean± SE, p**<0.01). (C) *twist* ISH of st24 embryos (arrowheads show normal migration, brackets impaired migration, asterisk the eye). Migration index and percentage of embryos with NC migration (n_CTLMO_=22, n_CX43MO_=24, n_Cx43MO+n-cad_=21, N=3, median ± quartile, mean ± SE p***<0.001). (D) Cell dispersion analysis after 6hrs of culture using delauney triangulation and its quantification (n=10, mean ± SE, p***<0.001, p**<0.01). (E) NCs expressing nRFP+mRFP (white arrows show normal protrusions, arrowheads short protrusions, green arrows N-cadGFP at cell membrane) and quantification of cell morphology (n_CTLMO_=51, n_Cx43MO_=32, n_Cx43MO+n-cadGFP_=58, median ± interquartile range, p***<0.001). (F) qPCR of n-cad expression (N=8, mean ± SE, p**<0.01). (G) *n-cad* ISH, arrowheads indicate normal and bracket irregular expression, white arrow injected side and green segmented line the NC. Quantification of *n-cad* expressing embryos (n_CTLMO_=68, n_CX43Mo_=61, N=9, mean ± SE, p***<0.001). (H) Rate of *n-cad* degradation (N=4, mean ± SE). See also Figure S2

Since our experiments suggest that Cx43 acts as regulator of NC migration via n-cad, we tested whether we could rescue the absence of Cx43 by restoring n-cad levels. Co-injection of Cx43MO with n-cad mRNA leads to a perfect rescue of the *in vivo* migration defects induced by the Cx43MO alone (Figure 3C). Moreover, the dispersion of NC explants, inhibited by Cx43MO, was rescued by co-injection of n-cad mRNA (Figure 3D; Supplemental Movie S5). Finally, when exogenous n-cad-GFP was delivered together with Cx43MO, we observed N-cad localisation at cell contacts, restored cell polarity and cell morphology that was similar to that found in CTLMO cells (Figure 3E). In conclusion, these results support the notion that Cx43 controls cell migration by regulating n-cad expression.

To determine whether Cx43 controls N-cad by regulating the levels of n-cad mRNA, we performed qPCR and ISH against n-cad mRNA. Both methods showed a significant reduction on n-cad mRNA in Cx43MO embryos (Figure 3F, G), suggesting that Cx43 regulates n-cad mRNA levels by either stabilising n-cad mRNA or promoting n-cad transcription. To investigate the first hypothesis, we measured mRNA stability by performing qPCR in the presence of the transcriptional inhibitor actinomycin D (Figure S2A-B). We found a similar rate of n-cad mRNA degradation between CTLMO and Cx43MO embryos (Figure 3H), indicating that Cx43 does not regulate mRNA stability, and arguing for a regulation of n-cad transcription.

Since Cx43 is present in two forms: the full length molecule (Cx43FL), which are building component of GJs, and the smaller carboxy terminal fragments (Cx43iso) that are not part of the GJ structures, we asked which of these two forms was responsible for the control of n-cad expression. One possibility is that the Cx43FL form regulates n-cad expression through GJ activity. In order to test this hypothesis, we measured N-cad expression in cells or embryos treated with two different GJ blockers. By analysing immunostaining in NC cultures, N-cad appeared unaltered in NC explants treated with flufenamic acid (FFA) or meclofenamic acid (MFA) in comparison to controls incubated with DMSO (Figure S2C). Furthermore, qPCR analysis showed that n-cad mRNA was unaffected by the use of GJ blockers (Figure S2D). These results suggest that the GJ activity is not involved in n-cad regulation, and open the possibility for a role of the Cx43iso as transcriptional regulators of n-cad.

### Cx43Tail controls n-cad transcriptional regulation and this activity is linked to its presence at cell nucleus

After discovering that GJ channels are not involved in the regulation of n-cad, we investigated other routes of Cx43 activity. To test which domain of Cx43 is involved in n-cad regulation, we co-injected Cx43MO with a full-length (Cx43FL), a truncated form missing the carboxy terminal (Cx43Trun), or a carboxy tail construct (Cx43Tail) (Figure 4A). Hence, the Cx43 form that is able to rescue the morphant phenotype contains important signaling involved in n-cad regulation. Co-injection of Cx43MO with Cx43FL or Cx43Tail mRNA rescued migration defects, showing a phenotype similar to the CTLMO group, while co-injection with the Cx43Trunc failed to rescue *in vivo* migration defects (Figure 4B). Moreover, similar to the effect on migration, n-cad deficiencies in Cx43MO embryos was rescued by the Cx43FL and Cx43Tail, but not by the Cx43Trunc mRNA (Figure 4C-4D; S3A). Importantly, when analysing cell polarity, we found that the inhibition on cell polarity induced by Cx43MO can be rescued by co-injection of Cx43FL and Cx43Tail, but not by Cx43Trunc (Figure 4E). Together, these results show that Cx43Tail is sufficient to promote migration by regulating n-cad levels and cell polarity.

**Figure 4.**
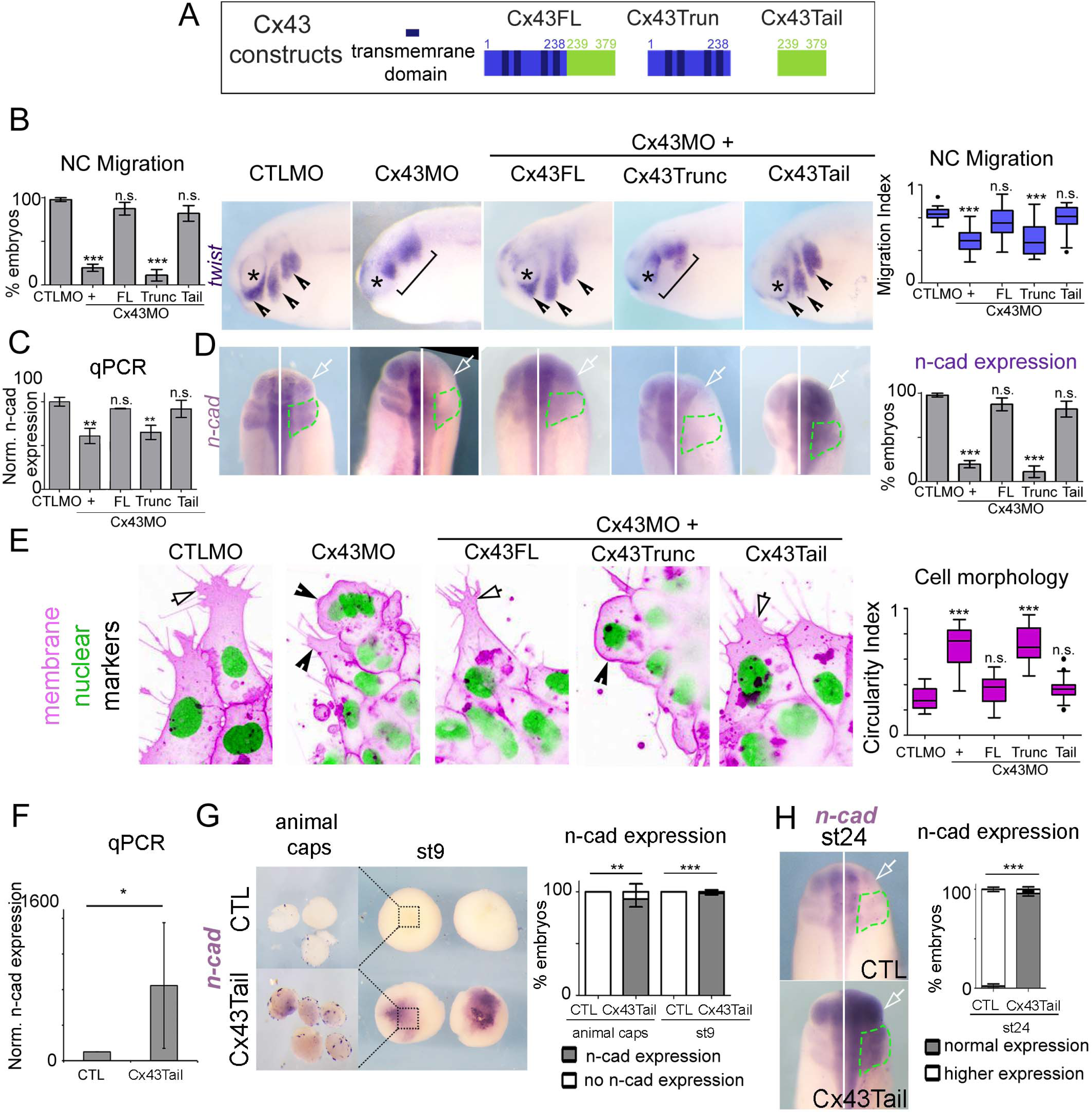
Cx43Tail is the n-cad transcriptional regulator. (A) Diagram of Cx43 constructs used. (B) Quantification of normal NC migration (% of embryos), *twist* ISH of st24 embryos (arrowheads display normal, bracket impaired migration, asterisk the eye) and NC migration index (n_CTLMO_=22, n_Cx43MO_=41, n_Cx43MO+FL_=21, n_Cx43MO+Trunc_=16, n_Cx43MO+Tail_=30, N=4, mean ± SE, median ± quartile, p***<0.001). (C) qPCR plotted as normalized n-cad expression (N=4, mean ± SE, p**<0.01). (D) *n-cad* ISH of st24 embryos, arrow shows injected side and green line the NC area. Quantification of *n-cad* expressing embryos (n_CTLMO_=42, n_Cx43MO_=42, n_Cx43MO+FL_=13, n_Cx43MO+Trunc_=15, n_Cx43MO+Tail_=19, N=4, mean ± SE, p***<0.001). (E) NCs with nuclear (green) and membrane probes (magenta), arrows indicate normal and arrowheads short protrusions. Circularity index (n_CTLMO_=20, n_Cx43MO_=32, n_Cx43MO+Cx43FL_=32, n_Cx43MO+Cx43Trunc_=23, n_Cx43MO+Cx43Tail_=26, median ± quartile, p***<0.001). (F) qPCR for relative n-cad expression of animal caps (N=4, mean ± SE, p*<0.05).(G) *n-cad* ISH of animal caps and st9 embryos, squares in the embryos indicate the dissected animal cap area. Quantification of *n-cad* expressing embryos (animal caps: n_CTLMO_=14, n_Cx43Tail_=45, st9 embryos: n_CTL_=22, n_Cx43Tail_=39, N=3, mean ± SE, p***<0.001, p**<0.01). (H) *n-cad* ISH of st24 embryos, arrow shows injected side and green line the NC area. Quantification of *n-cad* expressing embryos (st24 embryos: n_CTL_=23, n_Cx43Tail_=42, N=4, mean ± SE, p***<0.001). See also Figure S3.

To test whether Cx43Tail has the ability to induce n-cad expression, we used Cx43Tail in overexpression experiments in three different systems: in blastula embryo (when n-cad is not normally expressed), in blastula ectoderm (called animal caps, which do not express n-cad; Aybar et al., 2003; Chakrabarti et al., 1992), and in neurula embryos (which express n-cad). In all these systems we observed an upregulation of n-cad when overexpressing Cx43Tail (Figure 4G-4H; S3B). Moreover, qPCR analysis of Cx43Tail animal cap explants confirmed our results, also showing n-cad upregulation (Figure 4F). Taken together, these data show that the Cx43Tail is necessary and sufficient to regulate n-cad expression.

Cx43Tail could indirectly regulate n-cad expression, for example by modifying some signalling pathway, or it could work directly as a transcriptional regulator of n-cad. If Cx43 works as a transcriptional regulator, we would expect to find evidence of its localisation in the nucleus. We, therefore, investigated the subcellular localisation of Cx43 by expressing a low dose of Cx43Tail-GFP in the NC and observed a clear nuclear localisation of the tagged protein (Figure 5A). Similar nuclear localisation of Cx43Tail was obtained in mammalian Hela cells (Figure S4A) and amphibian XTC cells, using an antibody that reconginzes the carboxyl terminar (Figure S4F). To confirm these findings with endogenous Cx43 tail fragments (Cx43iso), we performed subcellular fractionation of *Xenopus* embryonic cells (Figure S5). Strikingly, Cx43iso was highly enriched at the nuclear fraction in comparison to all other fractions (Figure 5B).

**Figure 5.**
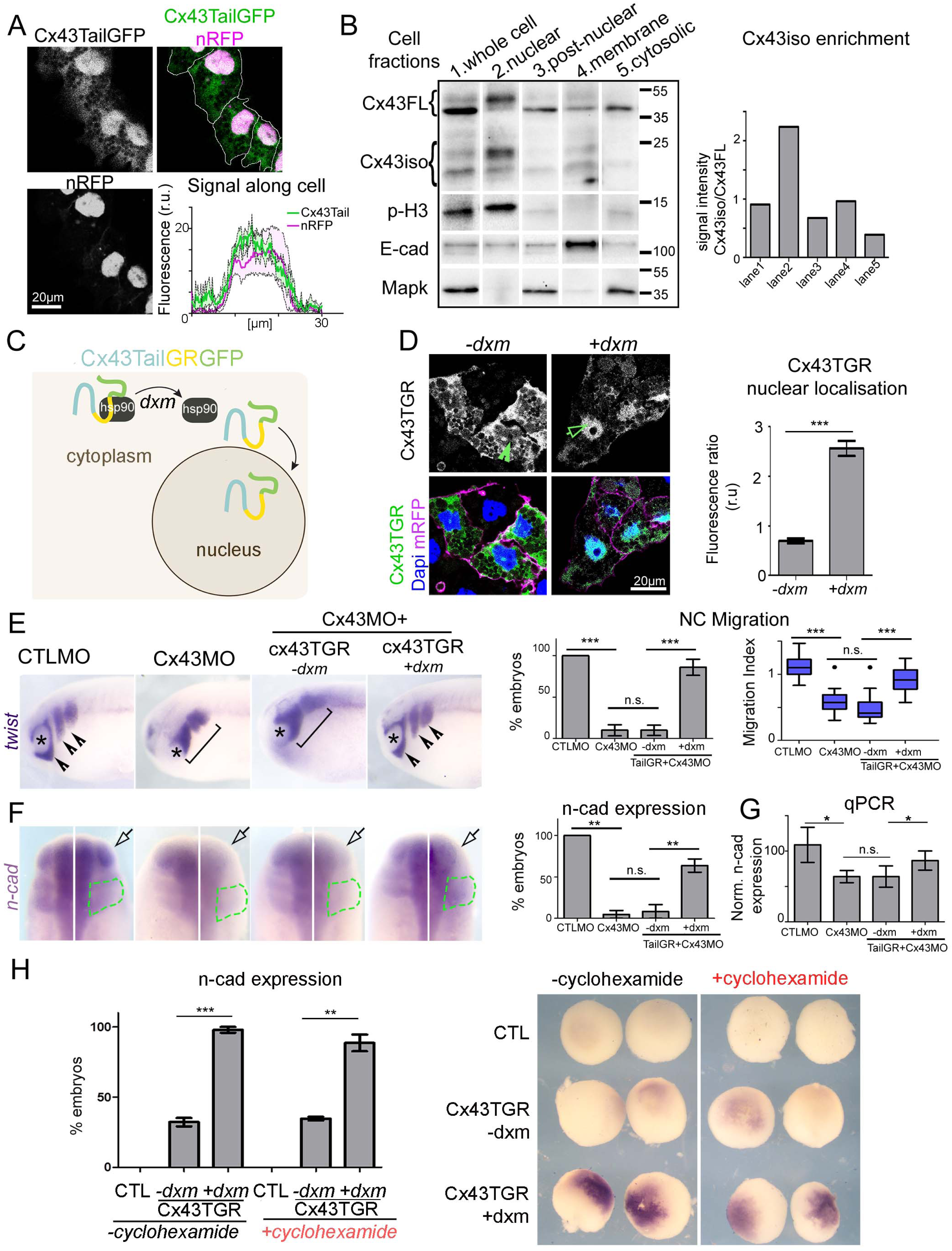
Cx43Tail nuclear localisation is requisite for its function. (A) Cx43Tail-GFP injected NCs and fluorescence along cell length (n=6, N=2). (B) Subscellular fractionation WB for Cx43, p-H3 (nuclear marker), E-cad (membrane marker) and Mapk (cytosolic marker). Cx43iso enrichment in proportion to Cx43FL. (C) Diagram of Cx43TGR-GFP function. (D) NCs expressing Cx43TGR-GFP +/-dxm. Cx43TGR nuclear fluorescence normalised to whole cell fluorescence (n_dmso_=23, n_dxm_=29, N=3, mean ± SE, ***p<0.001). (E) *twist* ISH of st23 embryos (arrowheads display normal migration, brackets impaired migration, asterisk the eye) and quantification of NC migration (n_CTLMO_=18, n_Cx43MO_=22, n_Cx43MO+Cx43TGR/dmso_= 18, n_Cx43MO+Cx43TGR/dxm_=19, N=4, mean ± SE, median ± interquartile range, p***<0.001). (F) *n-cad* ISH, white arrow shows injected side and green line the NC. Quantification of *n-cad* expressing embryos (n_CTL_=16, n_Cx43MO_=13, n_Cx43MO_+C_x43TGR/-dxm_=9, n_Cx43MO+Cx43TGR/+dxm_=13, N=3, mean ± SE, p**<0.01). (G) qPCR for relative n-cad expression (N=5, mean ± SE, p*<0.05). (H) Quantification of *n-cad* expressing embryos (-CHX: n_CTL_=35, n_Cx43TGR/-dxm_=41, n_Cx43TGR/+dxm_=28, +CHX: n_CTL_=27, n_Cx43TGR/-dxm_=23, n_Cx43TGR/+dxm_=25, N=3, mean ± SE, p***<0.001, p**<0.01). *n-cad* ISH for st9 embryos. See also Figures S4 and S5.

To investigate if the nuclear localisation of Cx43 is required for its transcriptional function, we used an inducible system that has been previously described (Tribulo et al., 2003). We fused the binding domain of the human glucocorticoid receptor (GR) with GFP tagged Cx43Tail (Cx43TGR-GFP, Figure 5C), allowing this construct to enter the nucleus only in the presence of dexamethasone (dxm). Indeed, dxm treated NCs displayed strong nuclear localisation of Cx43TGR (Figure 5D, +dxm), whereas its localisation was predominantly cytosolic in control NCs (Figure 5D, -dxm). These findings were mirrored in experiments performed on Hela cells (Figure S4D). To test the role of Cx43Tail nuclear localisation during migration, we used the inducible Cx43TGR system in NC migration and n-cad expression experiments. In both cases, we were able to rescue the morphant phenotype (reduced migration and decreased n-cad expression) when Cx43TGR was allowed to enter the nucleus using dxm treatment (Figure 5E-G). Similarly, in Hela or XTC cells, Cx43Tail overexpression led to elevated n-cad levels (Figure S4B, C, G), an effect similar to that found with the inducible system in Hela cells (Figure S5E). These data highlight the requirement for Cx43Tail in the nucleus to control n-cad transcription in different vertebrate systems, including *Xenopus* embryos and *Xenopus* and mammalian cell lines.

Our findings indicate that nuclear localisation of Cx43 is important for its function in promoting N-cad expression, yet it is not clear whether Cx43 regulates N-cad expression directly or whether it acts through an intermediary cascade of gene expresssion. To answer this question, we blocked all protein synthesis using the translation inhibitor cycloheximide (CHX), if Cx43 is not a direct regulator of N-cad transcription, it would require the transcription/translation of other proteins, and therefore any effect induced by Cx43 should be inhibited by CHX treatment. Cx43TGR-injected embryos treated with CHX, followed by dxm addition, showed an upregulation of n-cad expression, similar to non-CHX treated embryos (Figure 5H), therefore suggesting a direct effect of Cx43Tail on the n-cad transcriptional regulation.

### Cx43 requires BTF3 to function and localise at the nucleus

In order to further investigate the mode of direct regulation of Cx43 on n-cad, we performed mass spectrometry analysis of Hela cells transfected with Cx43Tail construct.

This provided a list of Cx43Tail binding partners that could be involved in n-cad regulation. Using 1.3-fold change between IP and control samples as the minimal limit for consideration (Gentzel et al., 2015), we found 30 candidates that are nuclear proteins and interact with Cx43Tail (Figure 6A). Examination of the Xenopus database, Xenbase, revealed that 18 of these candidates are expressed ubiquotisly in *Xenopus laevis* and only one is specifically expressed in migrating NC (Figure 6A): the basic transcription factor 3 (BTF3) (Cavallini et al., 1988; Kusumawidjaja et al., 2007; Zheng et al., 1990), which is enriched more than 1.49-fold in the IP sample in comparison to control.

**Figure 6.**
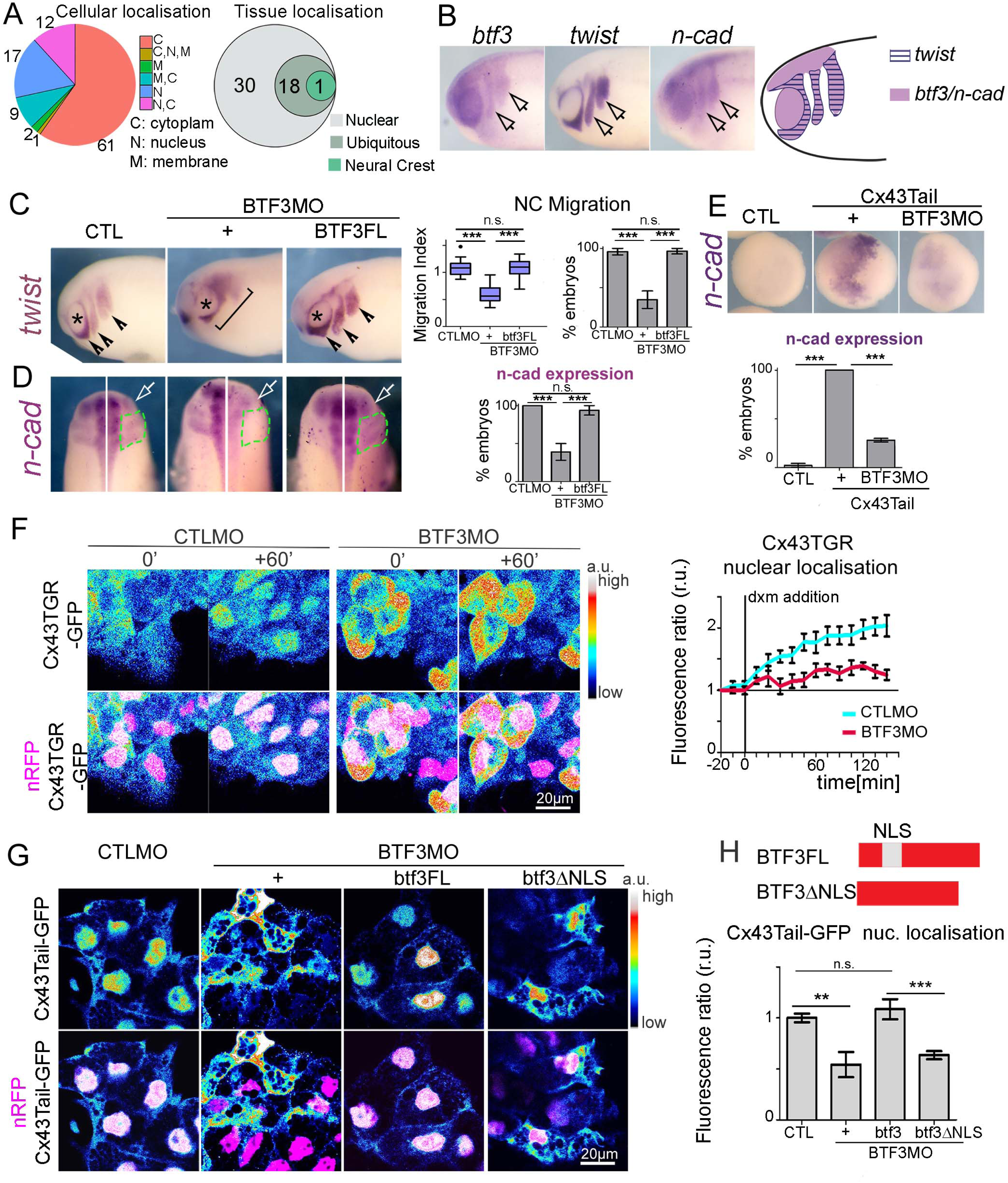
Cx43Tail requires BTF3 to function. (A) Summary of mass spectrometry results. Left: Pie chart of Cx43Tail interactants. Rigth: Venn diagram of interactants found in nucleus. (B) *btf3, twist* and *n-cad* ISH of st24 embryos, arrowheads show NC, arrows the neural tube and asterisk the eye. Summary of *btf3, twist* and *n-cad* profile. (C) *twist* ISH and quantification of NC migration (n_CTLMO_=19, n_BTF3MO_=36, n_BTf3MO+BTF3FL_=23, N=4, mean ± SE, p***<0.001, p**<0.01). (D) *n-cad* ISH. Quantification of *n-cad* expressing embryos (n_CTLMO_=14, n_BTF3MO_=35, n_BTf3MO+BTF3FL_=13, N=3, mean ± SE, p***<0.001). (E) *n-cad* ISH for st9 embryos. Quantification of *n-cad* expressing embryos (n_CTL_=24, n_Cx43Tail_=12, n_Cx43Tail+BTF3MO_=25, N=3, mean ± SE, p***<0.001, p**<0.01). (F) Cx43TGR-GFP NCs during dxm treatment. n-RFP is used to vislaize the nucleous. Heatmap of Cx43TGR-GFP intensity in NC. Cx43TGR-GFP nuclear fluorescence normalised to whole cell fluorescence (n=10, N=2, mean ± SE, p***<0.001). (G) Heatmap Cx43Tail-GFP intensity in NCs. (H) Diagram of BTF3 constructs. Quantification of Cx43Tail-GFP nuclear localisation (n_CTLMO_=17, n_BTF3MO_=8, n_BTF3MO+btf3FL_=12, n_BTF3MO+btf3dNLS_=15, N=3, mean ± SE, p***<0.001). See also Figure S6.

We validated *btf-3* expression pattern by ISH and confirmed that *btf3* is expressed at NC (Figure 6B), similar to *twist,* Cx43 and *n-cad,* with additional expression in eyes and neural folds matching the expression of *n-cad* and Cx43. This temporal and spatial co-localisation in the embryo and subcellular localisation in the nucleus suggests that Cx43Tail and BTF3 are potentially involved in the same pathway. To test whether BTF3 is involved in the same process as Cx43, we performed knock-down experiments using a BTF3 morpholino (BTF3MO), after confirming its efficiency (See star Methods, Figure S6A). Similar to Cx43MO, BTF3MO perturbed NC migration and n-cad expression (Figure 6C, D; S6C). Coinjection of BTF3MO with btf3FL that does not bind the MO, rescued impaired migration and n-cad expression (Figure 6C,D), thereby ensuring the specificity of BTF3MO. The effect of BTF3MO is specific for NC migration and does not affect NC induction, as shown by *snail2* ISH (Figure S6B). The similar phenotypes observed with BTF3 and Cx43 loss-of-function suggest that these molecules work together. Analysis of n-cad expression in blastula embryos showed that while embryos injected with Cx43Tail present a clear induction of n-cad expression, co-injection of BTF3MO with Cx43Tail significantly decreased this effect (Figure 6E), indicating that BTF3 is required for Cx43Tail activity.

Since Cx43 nuclear localisation is requisite for n-cad expression, we investigated the effect of BTF3MO on this process. For this purpose, we performed a temporal analysis of Cx43TGR nuclear localisation after addition of dxm in presence or absence of BTF3. Control NC showed a clear increase in the leves of Cx43Tail nuclear localization after dxm addition, while these levels remain lower in BTF3 depleted cells (Figure 6F), indicating that the nuclear localisation of Cx43Tail is BTF3 dependent. The converse, however, is not true: inhibition of Cx43 by Cx43MO does not affect the nuclear localisation of BTF3 (Figure S6D). These results suggest that BTF3 is localised in the nucleus by default, while Cx43 requires a partner to control its subcellular localisation. These observations are consistent with the presence of a nuclear localisation signal (NLS) in BTF3 (Wang et al., 2014) and its absence in Cx43Tail, and prompted us to test the role of the NLS of BTF3 on the nuclear localisation of Cx43Tail. As described above, nuclear localisation of Cx43 is impaired by BTF3MO but can be rescued by BTF3FL (Figure 6G). In contrast, the expression of a BTF3 construct lacking its NLS (Figure 6H) is unable to rescue Cx43Tail nuclear localisation (Figure 6G). Altogether these data indicate that one role of BTF3 is to recruit Cx43Tail into the nucleus by providing it with an NLS.

### Cx43tail and BTF3 directly interact and promote n-cad gene regulation

Co-localisation of BTF3 and Cx43Tail in the nucleus (Figure 7A) suggests interaction and a common role in that cell compartment, giving rise to the hypothesis that BTF3 interacts directly with Cx43. To test for physical interaction between Cx43 and BTF3, we used the bimolecular fluorescence complementation system (BiFC), in which two VENUS (variant of enhanced yellow fluorescent protein) components (VC and VN9m) form a fluorescent protein when are brought into close apposition (Saka et al., 2007). To build the BiFC system we fused the VENUS component (VC) to either Cx43Tail, Cx43Trunc, and the other VENUS component (VN9m) to BTF3, empty VC was used as a control. These constructs were co-injected into the NC along a nuclear marker, observing that Cx43TailVC co-injected with BTf3VN9m returned a positive signal, but not the other combinations (Figure 7B). Moreover, this signal was higher in the nucleus, where both molecules are typically found, therefore supporting our hypothesis of direct interaction between Cx43 and BTF3.

**Figure 7.**
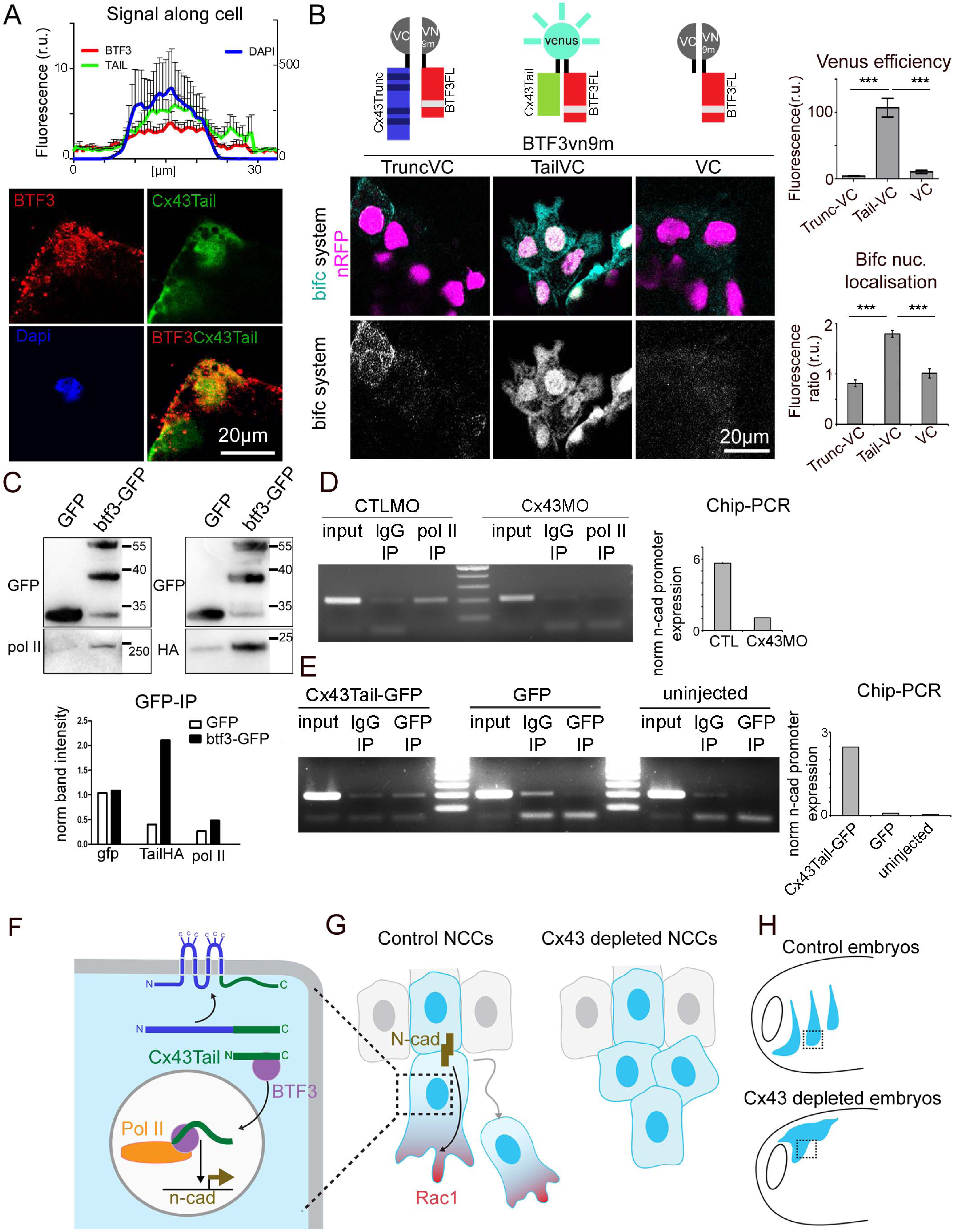
Cx43Tail and BTF3 form a transcriptional complex. (A) Fluorescence along cell length (n=9, N=2). Anti-BTF3, Cx43Tail-GFP and Dapi in NCs. (B) BiFC system. Diagram of constructs and function. NCs coexpressing VENUS components. VENUS efficiency and BiFC nuclear signal normalised to whole cell signal (n=16, N=3, mean ± SE). (C) WB of IP for GFP, Pol II, HA and normalised intensity graph. (D) ChIP-Pol II PCR for n-cad promoter. ChIP signal normalised to input (N=2). (E) ChIP-GFP PCR for n-cad promoter. ChIP signal normalised to input (N=2). (F) Cx43Tail (green), BTF3 (red), Pol II (brown) n-cad (black). (G) Control (cyan) and Cx43 depleted (orange) NCs, N-cad (black). (H) Embryos with CTLMO NC (light blue) and Cx43MO NC (orange). See also Figure S7.

To verify the biochemical interaction of Cx43Tail and BTF3, we performed immunoprecipitation (IP) using BTF3-GFP or GFP (as control) in conjunction with GFPTrap beads. We found that Cx43Tail-HA co-precipitates with GFP in BTF3-GFP expressing cells, but not in control GFP expressing cells (Figure 7C). Our experiments also confirmed that BFT3 interacts with Pol II (Figure 7C), as it has been previously shown (Zheng et al., 1987). These results support the notion that the three proteins, Cx43Tail, BTF3 and Pol II, interact and may function as a transcriptional complex.

To test this possibility, we performed chromatin immunoprecipitation (ChIP) using embryonic lysates and a Pol II antibody. We examined whether Pol II binds to the promoter of n-cad in presence or absence of Cx43, by performing PCR against the n-cad promoter region using CTLMO and Cx43MO ChIP samples as templates (see StarMethods). We detected that Pol ll is enriched at the n-cad promoter in CTLMO ChIP samples, whereas this enrichment was drastically reduced in ChIP samples collected from Cx43MO-injected embryos (Figure 7D). We conclude, therefore, that Cx43 is required for Pol II to bind at the n-cad promoter region so as to drive n-cad transcription.

To test whether Cx43Tail is directly recruited to the n-cad promoter we performed ChIP using Cx43TailGFP or Cyto-GFP (as a negative control) embryonic lysates and a GFP antibody. ChIP-PCR analysis showed that Cx43Tail is enriched at *n-cad* promoter region (Figure 7E). Therefore, we propose that Cx43Tail together with BTF3 and Pol II form a complex and control the transcriptional activity of n-cad.

## Discussion

Although Cx43 is mostly known for its function in GJs, some studies have suggested that it may also be involved in gene regulation independent of its GJ activity (Giepmans, 2006; Kang et al., 2009). These studies propose an indirect role for Cx43 in which it acts on other gene regulators, such as members of the CCN family (Fu et al., 2004; Gupta et al., 2001) or Smads (Takamatsu et al., 2007). Our findings identify a different mechanism, in which the Cx43Tail is directly involved in gene regulation. We found that Cx43Tail fragments are present in the NC of the developing embryo, as it has been shown in other cell types (Salat-Canela et al., 2014; Smyth and Shaw, 2013; Ul-Hussain et al., 2014). Using a combination of *in vivo* studies in the *Xenopus* embryo and experiments in other vertebrate cell systems, we characterized the physiological role of Cx43Tail signalling and uncovered a direct role in regulating n-cad expression. We further identified the molecular mechanism responsible for Cx43-driven n-cad regulation and showed how this activity is crucial during tissue morphogenesis.

Our findings provide important insight into this process and identify a previously unknown role for a protein primarily associated with cell-cell signalling. Specifically, we propose the following mechanism: Cx43Tail binds to BTF3 and, together, these proteins translocate into the nucleus where they form a complex with Pol II; this, binds to the n-cad promoter and activates n-cad transcription (Figure 7F). On the cellular level, once N-cad is transcribed and translated, it localises at cell contacts where it is known to modulate the polarization of Rac1 activity during CIL (Figure 7G; Theveneau et al., 2010; Scarpa et al., 2015). CIL in turn is required for the directional and collective migration of the NC, a process which is essential for morphogenesis (Figure 7H).

It is interesting to note that Cx43 has previously been shown to function as an adhesive molecule (Elias et al., 2007, 2010; Elzarrad et al., 2008), suggesting a direct role in controlling cell movement. Our findings highlight an alternative mechanism, independent of a direct involvement of Cx43 in cell adhesions. However, it is tempting to speculate that these functions may work in concert during cell migration.

In previous studies, it has been proposed that the generation of Cx43Tail isoforms is controlled by the PI3K/AKT/mTOR pathway in mammalian cells (Smyth and Shaw, 2013), the Mnk1/2 pathway in various cancer cell lines (Salat-Canela et al., 2014) and hypoxic conditions (Ul-Hussain et al., 2014). It is plausible that similar mechanisms are at play in our system. Indeed, in NC, the PI3K/AKT/mTOR pathway is present (Nones et al., 2012) and could, therefore, be implicated for the generation of Cx43Tail. It is also possible that Cx43Tail is produced in response to hypoxia and the Hif-1alpha pathway (hypoxia-inducible transcription factor-1), which are known to regulate NC migration (Barriga et al., 2013). Our study shows that the appearance and function of the Cx43 carboxy tail isoforms in the migrating NC coincide in time and space with the activity of Hif-1alpha, supporting findings in mammalian cells (Ul-Hussain et al., 2014).

Our results show that Cx43 control cell polarity, a finding that is consistent with known activities of Cx43 in cardiac NC (Xu, 2001; Xu et al., 2006). Cx43 has also been implicated in cell migration and polarity in other cell types (Elias et al., 2010; Homkajorn et al., 2010; Xu et al., 2006). In mouse embryonic fibroblasts this was attributed to the interaction of full-length Cx43 with microtubules (Francis et al., 2011). Although we found the cytoskeleton to be affected in Cx4MO NCCs, we observed that Cx43Tail alone has the ability to recover normal cell polarity and morphology. While we cannot exclude a direct effect of full-length Cx43 on microtubule organisation in our system, our findings show a different mechanism, in which Cx43 controls cell polarity and the cytoskeleton via the regulation of n-cad levels. Thus, we have established a direct link between Cx43 and contact-dependent cell polarity, which is likely to be conserved in other cellular system.

Cx43 appears to be at a convergent point of cell communication, cell polarity and n-cad regulation, but its effect on N-cad varies depending on context. Xu and colleagues have shown that Cx43 depletion led to N-cad decrease in cardiac NC in mice (Xu, 2001), while this was not the case in mouse neuronal cells derived from the developing cortex (Elias et al., 2007). This differential response could be explained by the presence of different pathways linked to Cx43 - one involving Cx43 full-length and another involving Cx43Tail. In this situation, factors controlling Cx43Tail production would define the composition of adhesion molecules at cell contacts, and this could be tuned differently in a time- and context-dependent manner.

We have identified a novel pathway by which an adherens junction component is regulated by a protein normally associated with a different cellular function, opening the possibility of cross-talk between processes mediating intercellular communication and cell-cell interaction and a possible mechanism of cross-regulation between gap and adherens junctions at the transcriptional level. Importantly, the high conservation of Cx43 and the presence of Cx43Tail in various vertebrate systems may reflect an evolutionary mechanism, where cells employ either one or the two pathways to control cell polarity during cell migration. In this study, we uncovered the Cx43Tail pathway, which displays a distinct function independent of the activities produced by Cx43 full-length, and provides a link between the regulation of N-cad expression and polarity defects. In the light of this new evidence, the role of Cx43 in collective migration during development or disease could be re-approached.

## Author Contributions

MK and RM conceived the study and designed experiments, with additional input from EHB. MK performed most experiments and data analysis. EHB helped with the bifc, promoter and Chip experiments. JL assisted with cloning. AS helped with cell line experiments; MG performed the MS analysis. MK and RM wrote the manuscript with input from all authors.

## Acknowledgements

We thank A. Streit (King’s College London) for comments on the manuscript. J. Smith (Crick’s Institute) for providing us with the bifc vectors and D. Becker (University College London) for help in the initial experiments on connexins. This study was supported by grants from MRC (J000655 and M010465), BBSRC (M008517) and Wellcome Trust to RM. MK was supported by a Latsis Postgraduate Scholarship, a DAAD short term research grant and travel grants from Boehringer Ingelheim Funds and the Company of Biologists. EHB was supported by fellowships from EMBO LTF-971 and a Marie Skłodowska Curie IF-2014_ST 658536.

## Supplemental Figures

**Figure S1.**
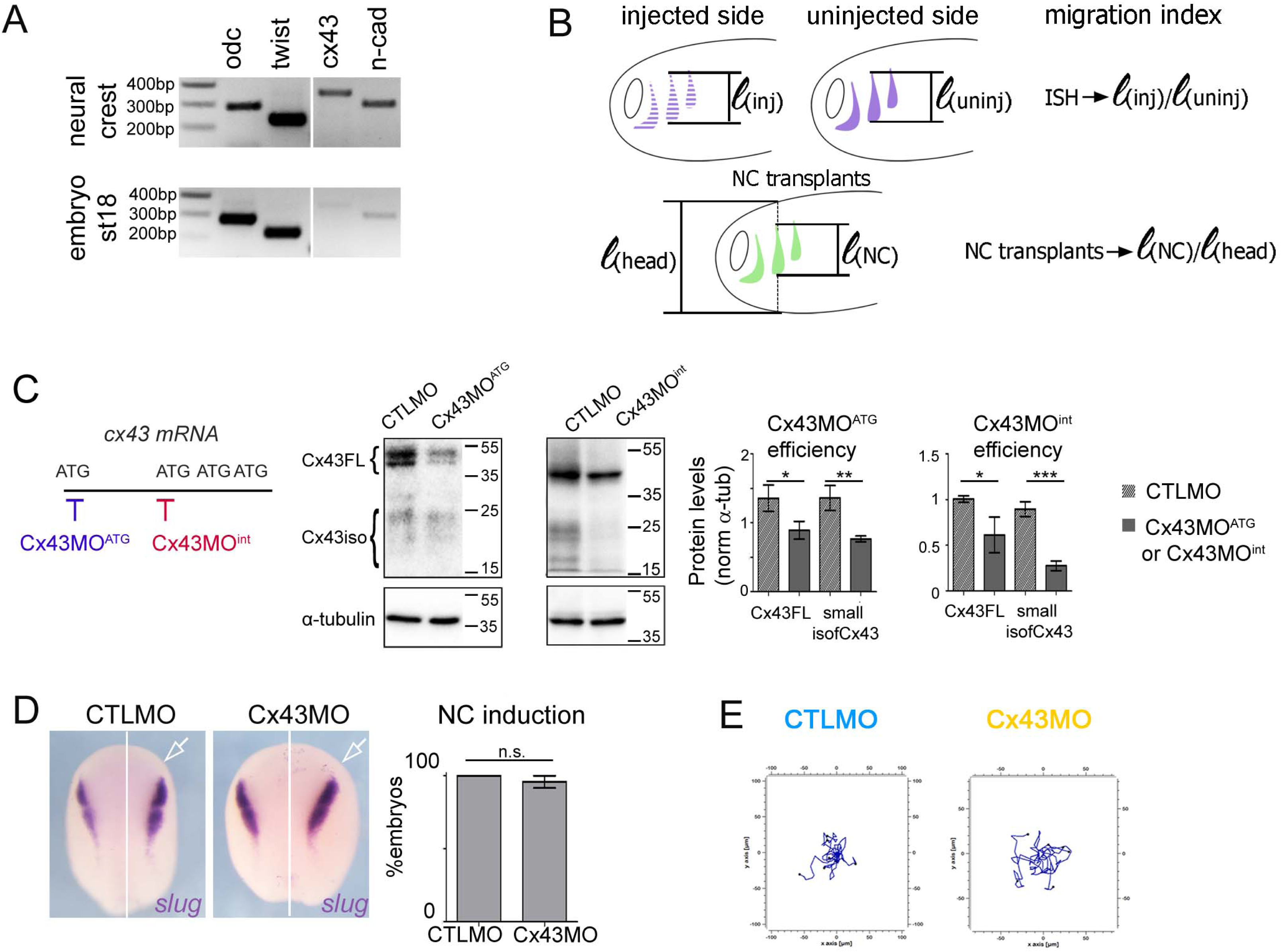
Related to Figure 1. (A) RT-PCR for NC dissected from st18 embryos and for st18 whole embryos for: ubiquitous marker odc, neural crest marker twist, cx43 and n-cad. (B) Diagram of migration index analysis for ISH and NC transplants, 𝓵: measured length. (C) Diagram of MO targets and WB for Cx43 and α-tubulin. Cx43 levels normalised to α-tubulin, (N=3, mean ± SE, p*<0.05, p**<0.01, p***<0.001). (D) *snail2ISH* for stage 16 embryos, arrow indicates the injected side. Quantification for NC induction (n_CTLMO_ =20, n_Cx43MO_ =15, N=4, mean ± SE). (E) Representative motility single cells tracks.

**Figure S2.**
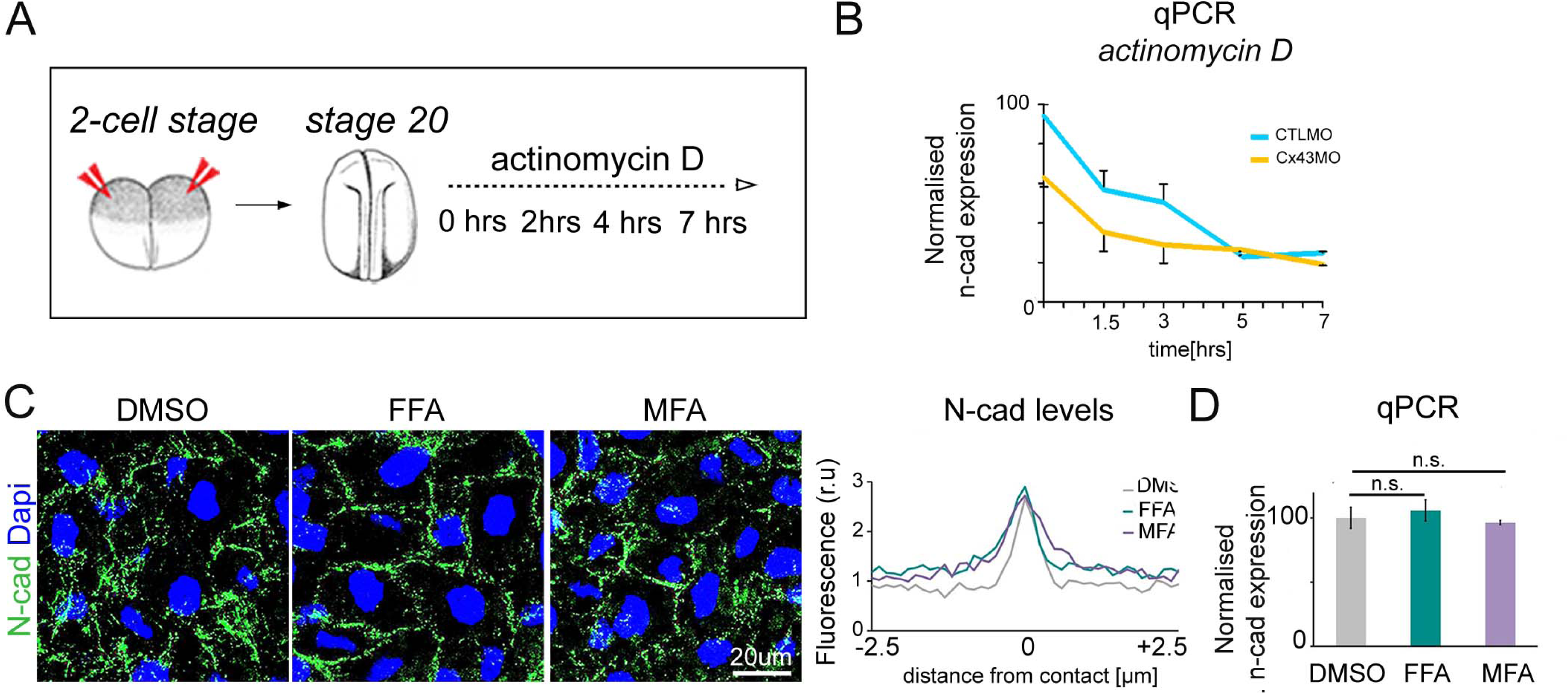
Related to Figure 3. (A) Summary of the actinomycin D treatment. MO injection at the 2-cell stage, followed by actinomycin D treatment at stage 20, and collection of samples at the indicated times. (B) qPCR plotted as relative n-cad expression over time with 4μg/ml actinomycinD treatment (N=4). (C) Immunostaining for N-cad using dmso or GJ blockers FFA and MFA. N-cad levels, 0 represents contact point (n=15, N=3, mean ± SE). (D) qPCR plotted as a relative n-cad expression (N=3, mean ± SE).

**Figure S3.**
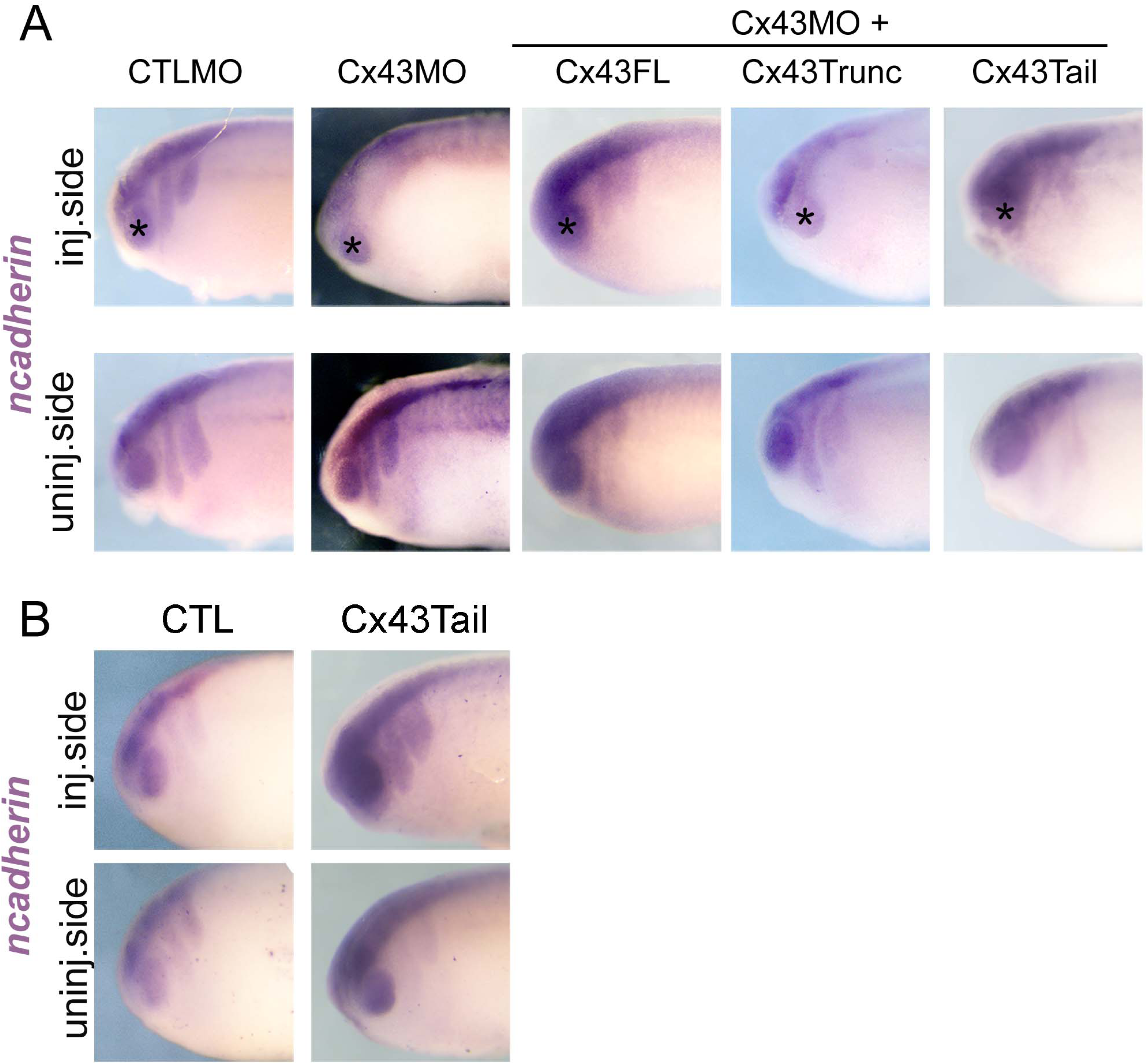
Related to Figure 4. (A) Lateral view of embryos showing *n-cad* ISH of the same embryos shown in dorsal view in Figure 4D and (B) from Figure 4H. The injected and injected side of the same embryo are shown for comparison.

**Figure S4.**
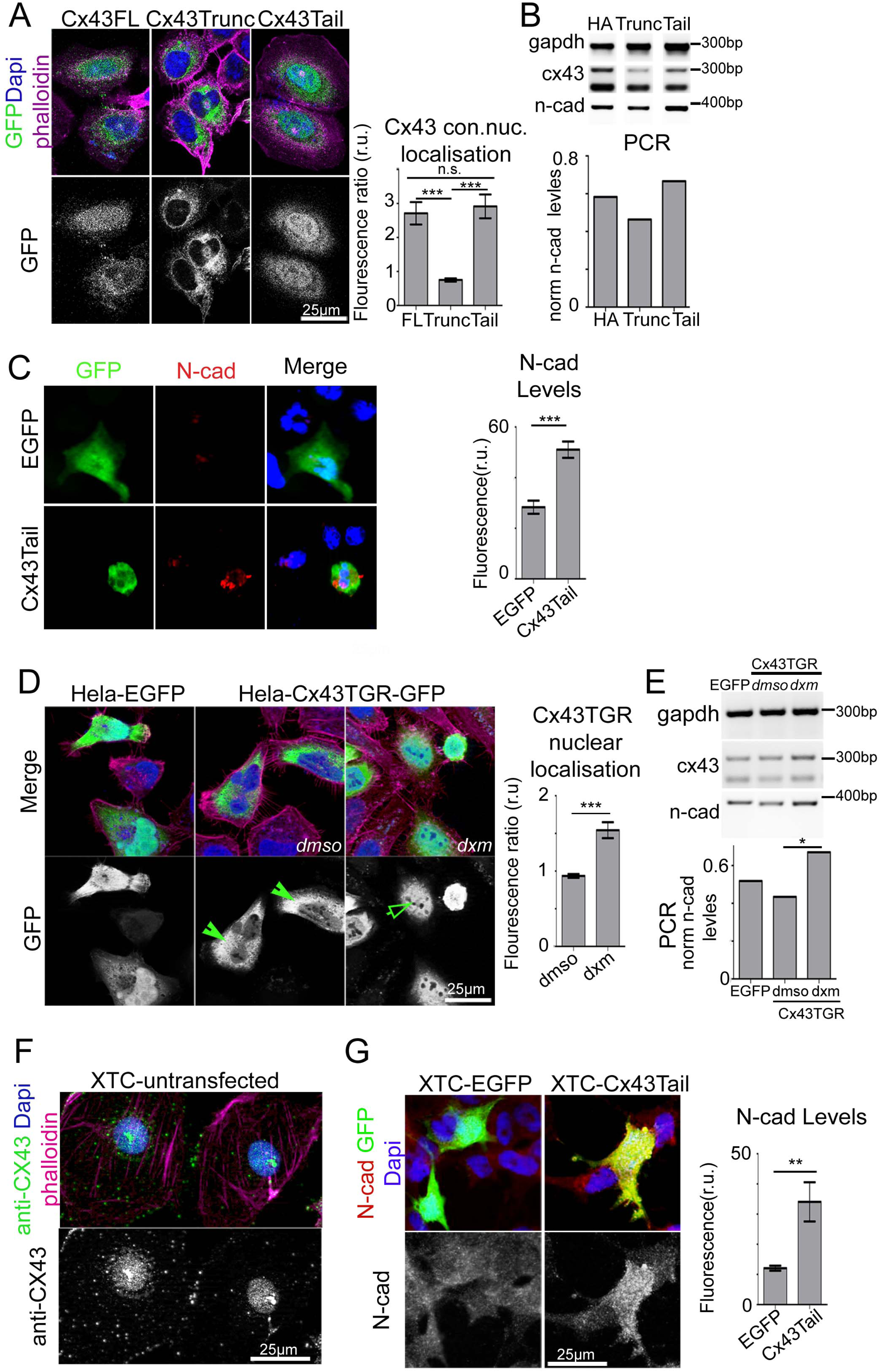
Related to Figure 5. (A) Hela cells transfected with Cx43 constructs tagged with GFP (green). GFP nuclear fluorescence normalised to whole cell fluorescence (n=8, N=2, mean ± SE, p***<0.001). (B) PCR of Hela for Cx43 constructs using gapdh, cx43 and n-cad primers. PCR graph showing n-cad levels normalised to gapdh. (C) Immunostaining of Hela cells against N-cad. Quantification of N-cad levels (n_EGFP_=20, n_Cx43Tail_=15, mean ± SE, p***<0.001, p**<0.01). (D) Hela-EGFP and Hela-Cx43TGR-GFP cells treated with dmso or dxm. Cx43TGR-GFP nuclear fluorescence normalised to whole cell fluorescence (n_Cx43TGR/dmSo_=15, n_Cx43TGR/dmso_ =12, N=2, mean ± SE, p***<0.001). (E) PCR of Hela using gapdh, cx43 and n-cad primers. PCR graph showing n-cad levels normalised to gapdh. (F) Immunostaining of untrasfected XTC cells for Cx43. (G) Immunostaining of XTC cells against N-cad. Quantification of N-cad levels (n_EGFP_=17, n_Cx43Tail_=17, mean ± SE, p**<0.01).

**Figure S5.**
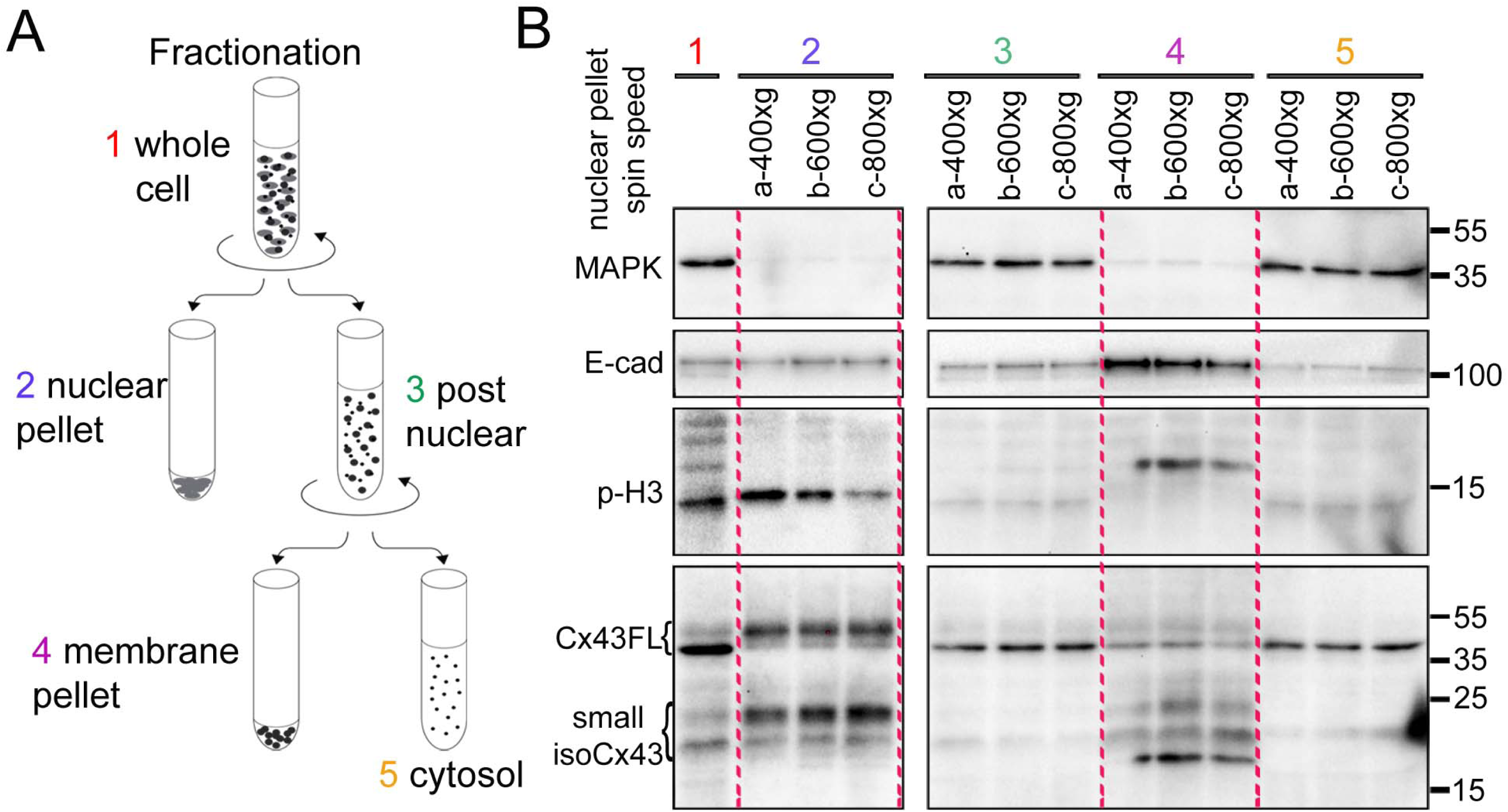
Related to Figure 5. (A) Diagram of subcellular fractionation (B) WB showing subcellular fractionation (see Figure 5B) of wild type *X*. embryos using (a) 400xg, (b) 600xg, (c) 800xg as the initial spin speed for nuclear pellets and Cx43, phopho-histone H3 (p-H3, nuclear control), E-cadherin (E-cad, membrane control) or Mapk (cytosolic control).

**Figure S6.**
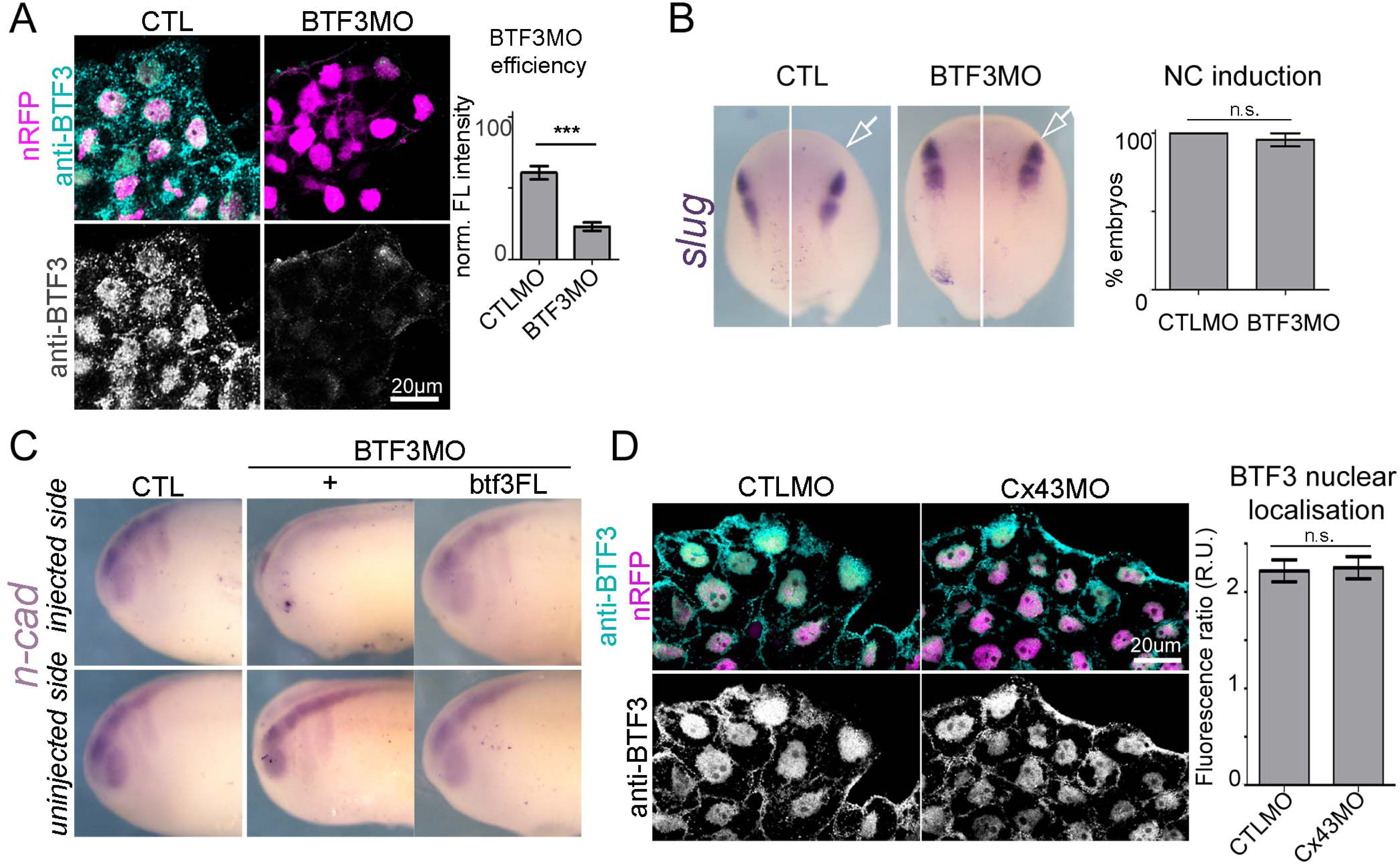
Related to Figure 6. (A) Immonostaining for BTF3. Fluorescence levels of BTF3 normalised to background levels (n_CTLMO_=23, n_BTF3MO_=27, N=3, mean ± SE, p***<0.001, p**<0.01). (B) *snail2* ISH for stage 16 X. laevis embryos, arrow indicates the injected side. Quantification of NC induction (n_CTLMO_ =15, n_BTF3MO_=10, N=2, mean ± SE). (C) Lateral view of *n-cad* ISH from embryos from Figure 6E. (D) Immunostaining for BTF3. BTF3 nuclear fluorescence normalised to whole cell fluorescence (n_CTLMO_=20, n_Cx43MO_=20, N=2, mean ± SE).

**Figure S7.**
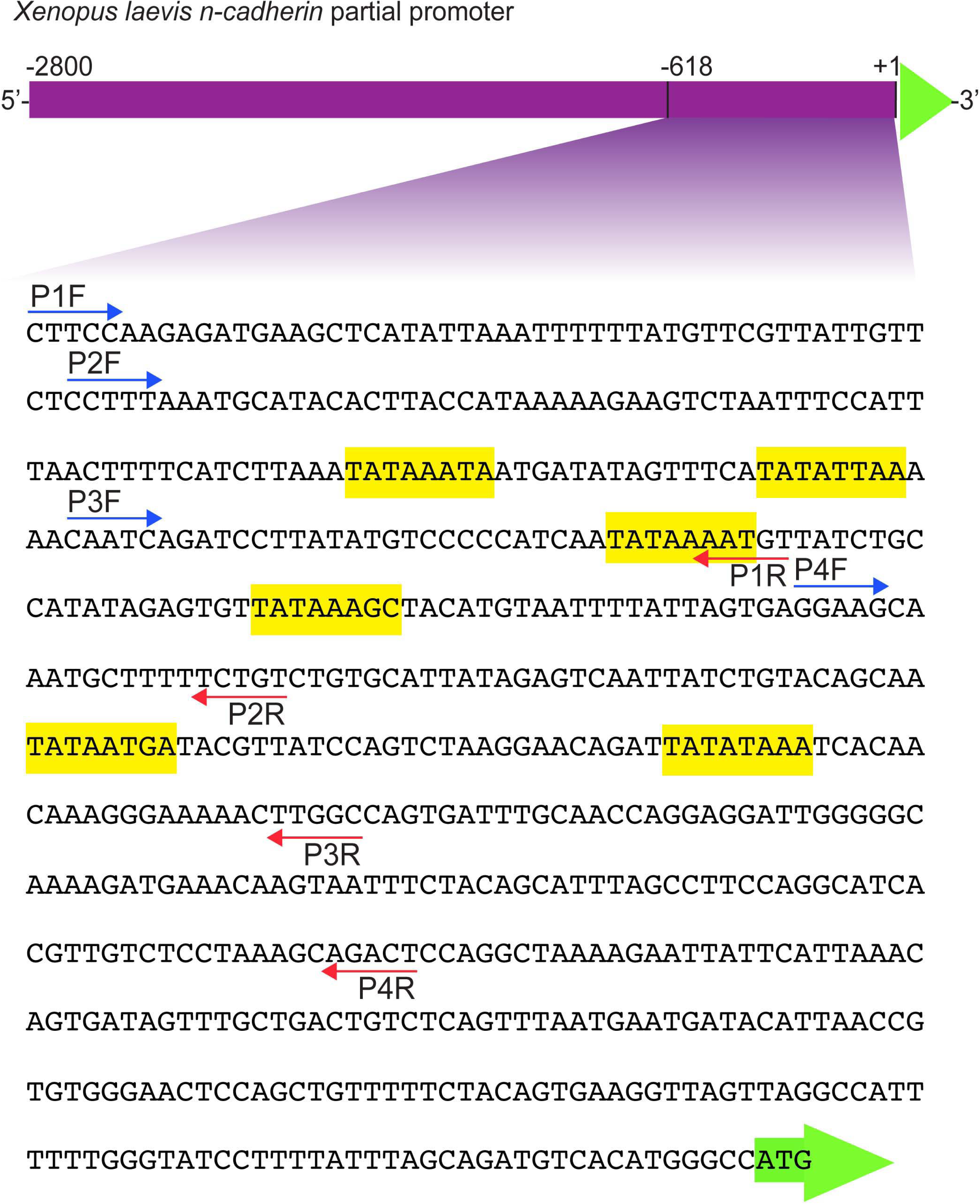
Related to Figure 7. Putative TATA boxes on partial n-cad promoter region form *Xenopus laevis* obtained using ElemeNT Analysis Resource (highlited in yellow) and binding sites for the primers used in the ChIP experiments.

## STAR Methods

### Embryo manipulation

Adult *Xenopus laevis* were maintained at 17°C used according to the regulations of the Animals (Scientific Procedures) Act 1986 as described previously (Steventon et al., 2009). *Xenopus laevis* embryos were obtained and staged as described previously (Newport and Kirschner, 1982; Nieuwkoop P., 1994). To specifically target the neural crest, embryos were injected into the animal ventral blastomeres at the 8- or 16-cell stage on one side for all experiments, except when embryonic lysates were used. In this case embryos were injected in both animal sides of 2-cell stage embryos. Embryo microinjections were performed according to (Aybar et al., 2003). If required, fluorescein-dextran (FDx; Invitrogen, D1821, 20 ng) or rhodamine-dextran (RDx; Invitrogen, D1824, 20 ng) were used as tracers. Oligomorpholinos against *Xenopus laevis* Cx43 (Cx43^ATG^MO; 24ng, 5'- TTCCTAAGGCACTCCAGTCACCCAT-3', Cx43^int^MO; 8ng, 5'- AAAAATGGTTTTCTTGTGGGTCGA-3') and BTF3 (BTF3MO, 40ng, 5'- AACGGACCGGGTTTAAAGGCTTCCT-3') were synthesised and provided by Gene Tools LLC. Equimolar concentrations of standard control morpholino (CTLMO: 3'- ATATTTAACATTGACTCCATTCTCC - 5') were used. Cx43^ATG^MO targets the region around the ATG of the full-length cx43 mRNA and Cx43^int^MO targets the region before the ATGs that guide the translation of the small cx43 fragments (Cx43MOint; Figure S1C). To test the efficiency of these MOs on the endogenous cx43 mRNA, we performed western blot analysis. Both Cx43MOs efficiently lowered the protein levels of the endogenous Cx43FL and Cx43iso (Figure S1C). In order to validate BTF3MO efficiency, we tested by immunostaining of NCs the BTF3 protein expression, which showed decreased levels of BTF3 in BTF3MO NCs compared to CTLMO (Figure S6A).

*In situ* hybridisation was performed as previously described (Fawcett and Klymkowsky, 2004; Harland, 1991). Digoxigenin-labelled RNA probes were prepared for *snail2* (Mayor et al., 1995), *twist* (Hopwood et al., 1989), *n-cadherin* (Theveneau et al., 2010) and *btf3* (Xenbase: XL075i22). *snail2* (0.8 μg/ml) was used to assess NC induction, *twist (0.7* μg/ml) for NC migration, *n-cadherin* (1 μg/mL) to assess expression levels and *btf3* (0.7 μg/mL) expression pattern. NC induction, NC migration and n-cad expression were plotted as a percentage of embryos for each of these process. NC migration was further assessed using migration index, which is defined as ratio of NC length of the injected side to that of the uninjected side or, in the case of NC transplants, as the ratio of NC length of the injected side to the dorso-ventral lenght of the head.

Whole mount immunostaining was carried out as previously described (Dent et al., 1989) using Cx43 antibody (1:1000, Sigma, C6219).

Animal cap explants were dissected from *Xenopus* blastulas (stage 8) using a standard technique (Aybar et al., 2003; Chakrabarti et al., 1992) and prepared for in situ-hybridisation as described above. For animal cap assays, mRNA was injected into the animal side of the two bastomeres of two-cell stage embryos.

### NC manipulation and imaging

Neural crest transplants were carried out as previously reported (De Calisto et al., 2005). For *in vitro* experiments, cranial NC explants were dissected at stage 18 using a standard technique (Alfandari et al., 2003; Borchers et al., 2000) and plated on fibronectin (Sigma, F1141) coated dishes as described previously (Theveneau et al., 2010). For single cell assays, NCCs were briefly dissociated in Ca^2+/^Mg^2+^-free Danilchick medium (Theveneau et al., 2010).

**Immunostaining** was performed on NC explants as previously described (Moore et al., 2013) using anti-N-cadherin (1:50, rat IgG, clone MNCD2, DSHB), anti-BTF3 (1:100, Abcam, ab107213), anti-rabbit IgG Alexa488 (1:500, Invitrogen, A11034) and anti-rat IgG Alexa488 (1:500, LifeTechnologies). When required, 4',6-diamidino-2-phenylindole (DAPI; 1 μg/mL, Sigma, D9542) and/or phalloidin tetramethylrhodamin-b-isothiocyanate (PhR; 1 μg/mL, Sigma, P1951) were used.

**Imaging of fixed embryos** was performed using a MZFLIII Leica fluorescence stereomicroscope equipped with a DFC420 Leica camera controlled by Leica IM50 software. **Imaging of *in vitro* neural crest migration** was performed using time-lapse cinematography as described previously (Carmona-Fontaine et al., 2008; Theveneau et al., 2010). For cell motility and migration, compound microscopes (either an Eclipse 80i Nikon microscope with a Hamamatsu Digital camera or a DMRXA2 Leica microscope with a Hamamatsu Digital camera controlled by Simple PCI program) equipped with motorized stages and a 10x/0.30NA dry lens were used. NC cultures were prepared as described above. Images were acquired every 3-5 min for a period of 10-12 hr. For cell morphology imaging, a TCS SP8 microscope with 20X/0.50NA or with a 63X/0.90NA water immersion objective lenses controlled by LAS-AF software was used. **Fixed cells** were imaged using a 63X/1.4NA oil immersion objective and a Leica TCS SPE confocal microscope controlled by LAS-AF software.

### Cell migration cell morphology analysis and CIL

Chemotaxis assay was performed following a standard procedure (Theveneau and Mayor, 2011) using heparin acrylic beads (Sigma, H5263) coated with 1 μg/mL purified human stromal cell-derived factor-1 (Sigma, SRP3276). Cell motility and chemotaxis were analysed using ImageJ (https://imagej.nih.gov) analysis tools, as described previously (Carmona-Fontaine et al., 2008; Matthews et al., 2008; Theveneau et al., 2010). Cell dispersion was analysed using Delaunay triangulation algorithm, which connects each cell nucleus with its closest neighbours (other cell nuclei) in such a way that a network of triangles covers the entire explant and plotted as average explant triangle area (Carmona-Fontaine et al., 2008). CIL was assessed for individually migrating cells by counting cell separation events 30 min after contact initiation (Scarpa et al., 2015; Theveneau et al., 2013) and collision angles (Carmona-Fontaine et al., 2008). Also, CIL was assessed for cell collectives by explant confrontation assay using the overlap index which is defined as the ratio of overlap area of two explants to total area of smaller explant (Kashef et al., 2009). Cell morphology was assessed by deploying the circularity index (complete circle=1) and estimated by ImageJ analysis tools.

### Molecular biology, plasmids, and reagents

For cDNA synthesis, total RNA was isolated from stage 23-24 *Xenopus* embryos as previously described (Barriga et al., 2013). Quantitative PCR (qPCR) was performed on an Applied Biosystems ABI 7900HT machine using the Fast SYBR Green Master Mix (Applied Biosystems, 4385612) and the following primers: ef-1 forward 5'-ACCCTCCTCTTGGTCGTTT-3', ef-1 reverse 5'- TTTGGTTTTCGCTGCTTTCT-3' (Hong and Saint-Jeannet, 2007), ncad forward 5'-CAGGGACCAGTTGAAGCACT-3', ncad reverse 5'-TGCCGTGGCCTTAAAGTTAT-3' (Nandadasa et al., 2009). n-cad mRNA expression was plotted as relative expression normalised against the housekeeping gene *ef-1.*

**Plasmids:** Cx43 constructs were synthesised using the *X*. *laevis* Cx43 sequence from cDNA clone (UniGene ID <u>XL.1109</u>) as template. Full length Cx43 (Cx43FL, aa 1-379), Cx43 carboxy terminally truncated construct (Cx43Trunc, aa 1-238) and Cx43 caboxy tail construct (Cx43Tail, aa 239-379) were cloned into 5’BamHI/3’XhoI of pCS2+ or pCS2-EGFP vectors. The pCS2-EGFP vector was kindly provided by Dr. Masa Tada. Inducible construct of Cx43Tail was prepared by fusing the Cx43Tail (aa 239-379) to the ligand-binding domain of the human glucocorticoid receptor (GR, aa 512-777). Cx43Tail was cloned into 5’EcoRI/3’SacI and GR into 5’SacI/3’XhoI of pCS2+ and pCS2-EGFP. BTF3 constructs were synthesized using the *X*. *laevis* sequence from cDNA clone (UniGene ID XL.3536). BTF3FL (aa 1-162) was cloned into 5’EcoRI/3’XhoI of pCS2+ or pCS2-EGFP. BTF3 deletion construct lacking the NLS region RRKKK (BTF3-dNLS, aa 1-158) was cloned into 5’BamHI/3’ClaI of pCS2+. For the BiFC experiments (Figure 7B), Cx43Tail and Cx43Trunc were cloned into 5’BamHI/3’BamHI of pCS2-VC155. BTF3FL was cloned into 5’BamHI/3’BamHI of pCS2-VN9m. BiFC vectors were kindly provided by Prof James C Smith. All construct sequences are verified by automated DNA sequencing (Source Biosciences, UK). Pasmids were linearised and mRNA transcribed as described before (Harland and Weintraub, 1985). The mRNA constructs injected were: membrane GFP (mGFP, 300 pg), membraneRFP (mRFP, 300pg) nuclearRFP (nRFP, 300 pg), lifeactin-GFP (400pg), N-cadherin-GFP (500pg), Cx43FL (500pg), Cx43Trunc (500pg), Cx43Tail (500pg), BTF3FL (500pg), BTF3-dNLS (500pg), Cx43Tail-VC (500pg), Cx43Trunc-VC (500pg), BTF3-VN9m (500pg). GFP-Lpd (Krause et al., 2004) was injected as DNA (800pg). All analysis on fluorescent constructs was assessed by normalizing to the background fluorescence and when required, to total cell area fluorescence.

The following **reagents** were used: flufenamic acid (50-100μΜ, Sigma, F9005), meclofenamic acid (50-100μΜ, Sigma, M4531), actinomycin D (20μΜ, Sigma, A1410), cycloheximide (10 μM, Sigma, C7698) and ethanol-dissolved dexamethasone (10μM) was added to the culture medium at stages 14-15 and maintained until NC migratory embryonic stages (stage 23). To control the possible leakage of inducible chimeras, a sibling batch of embryos were cultured without dexamethasone and processed for in situ hybridisation.

### Immunoprecipitation, Fractionation, and Western blotting

Whole embryos were prepared for western blot after homogenisation in lysis buffer (20 mM Tris, 100 mM NaCl, 0.01% Triton-X, pH 8.0) with added protease inhibitors (Roche, 11836153001) and phosphatase inhibitors (Roche, 04906837001) as described previously (Moore et al., 2013; Scarpa et al., 2015). For western blots, the following antibodies were used: Connexin43 (Sigma, C6219, 1:1000), N-cadherin (DSHB, clone MNCD2, 1:800), p42/44 MAPK (Cell Signalling, 9102S, 1:2000), α-tubulin (DSHB, clone 12G10, 1:1000), phospho-histone H3▭(Millipore, 06579, 1:2000), GFP ▭(Invitrogen, A11122, 1: 2000), rabbit IgG HRP-linked (Amersham ECL, NA934, 1:3000), mouse IgG HRP-linked (Amersham ECL, NA931, 1:3000) and rat IgG HRP-linked (Sigma, A9037, 1:2000). **Immunoprecipitation** was performed using GFP-TRAP beads (Chromotek) as previously described (Scarpa et al., 2015). **Fractionation** of *Xenopus laevis* embryos was performed using differential centrifugation protocols (Dimauro et al., 2012; Kushima et al., 2011) with modifications. Briefly, stage 18 *Xenopus laevis* embryo lysates using a 22G needle and 20 μL homogenisation buffer (HB: 250 mM sucrose, 20 mM Hepes, 10mM KCl, 1 mM EDTA, 1 mM EGTA, 1 mM DTT, phosphatase and protease inhibitors) per embryo. All samples were centrifuged at 250xg for 5 min to remove embryo debris and yolk. After collecting the supernatant, it was distributed to three different tubes and spun at three different speeds (400xg, 600xg or 800xg) for 5 minutes to isolate the cell nuclei. The post-nuclear supernatants were kept for further processing. The nuclear pellets were washed once more in buffer HG and centrifuged at the appropriate speed (400xg, 600xg or 800xg). Nuclear pellets were then resuspended in 40 μL 10% glycerol/ 0.1% SDS/ 1% Triton-X in HB. The post-nuclear supernatants were then centrifuged at 16,000 G for 30 min to isolate the membrane fraction. The supernatants were collected as the cytosolic fraction and the pellets as the membrane fraction. 40 μL of 10% glycerol/ 0.1% SDS/ 1% Triton-X in HB were added to the membrane pellet. Appropriate amount of sample buffer was added to each sample, and samples were processed for acrylamide gel electrophoresis and western blot analysis.

Western blot data were analysed using densitometric analysis tools from the Image Lab software. The average relative density of each protein signal (band on the blot) was normalised to the equivalent loading control (i.e. Mapk or α-tubulin). In order to avoid loading errors, the same membrane was blotted against the loading control after stripping as described before (Scarpa et al., 2015).

### Cell cultures

HeLa cells (Leibniz Institute Collections of Microorganisms and Cell Culture, DSMZ, Germany) were cultured in DMEM supplemented with 10% fetal calf serum (Life Technologies, CA, USA) at 37 °C in a humidified atmosphere of 10% CO_2_ and transfected as indicated using Rotifect (Carl Roth, Germany) according to the manufacturer’s instructions.

*Xenopus* embryonic fibroblasts (XTC, a kind gift from Ana Losada) were cultured in 67% DMEM/H_2_O supplemented with 10% fetal calf serum at 25°C in a humidified atmosphere of 5% CO_2_ and transfected as indicated using Viafect (Promega, USA).

Plasmids for transfection were as indicated.

Cells were fixed in 4% paraformaldehyde, immunostained against N-cadherin using MNCD2 monoclonal antibody (DSHB; MNCD2 was deposited to the DSHB by Takeichi, M. / Matsunami, H. (DSHB Hybridoma Product MNCD2) and imaged on a Zeiss Apotome (Zeiss, Germany).

### Mass spectrometry

Co-immunoprecipitation of the FLAG-tagged Cx43Tail for mass spectrometric analysis was performed as described previously (Gentzel et al., 2015). Subsequently of acid elution with 0.1 M glycine pH 2.5, the pH of the eluates was adjusted to pH 8.0 with 0.5 M Tris. For protein digestion samples were incubated overnight with 2 μg trypsin (Trypsin Gold, Promega, USA) followed by addition of 0.1 μg Lys-C (Roche, Germany) and incubation for additional 6h. The peptides were desalted on C-18 reversed phase stage tips (Nest Group, USA), dried in vacuum and stored at -20°C until analysis. Dried peptides mixtures were recovered in 3 μl 30% formic acid and diluted with water to 23 μl. Five pl of the digest were injected on a Nano-LC system (Eksigent425 2D Nano-LC; Sciex, USA) and peptides were separated by reversed phase chromatography on C-18 columns prepared in-house (Reprosil-Pur C18-AQ, 1.9 μm, Dr. Maisch, Germany, PicoFrit Columns, 75 μm i.d., New Obejctive, USA) in a linear gradient of 100% eluent A (0.1% formic acid) to 55% eluent B (60% actenitrile, 0.1% formic acid) in 120min. The LC-system was setup as vented column and sample was load and desalted at a flow rate of 400 nl/min and separation was carried out at a flow rate of 200 nl/min. The nano-LC system was hyphenated to the mass spectrometer (Q-Exactive HF, ThermoScientific, Germany) which was operated in data-dependent mode. Data interpretation was performed with Mascot V2.2 (Matrixscience, UK) and Progenesis LC-MS V4.1 (Nonlinear Dynamics, UK).

### Statistical Analysis

Comparison of percentages was performed using contingency tables as described previously (Taillard et al., 2008). Normality of data sets was tested using Kolmogorov-Smirnov’s test, d’Agostino and Pearson’s test and Shapiro-Wilk’s test using Prism6 (GraphPad). Datasets following normal distribution were compared with Student's t-test (two tailed, unequal variances) or ANOVA with a Dunnett's multiple comparisons post-test using Excel or Prism6 (GraphPad). Datasets that did not follow a normal distribution were compared using Mann Whitney’s test or a nonparametric ANOVA (Kruskal Wallis with Dunn’s multiple comparisons post-test) using Excel or Prism6. Cross-comparisons were performed only if the overall *P*-value of the ANOVA was less than 0.05.

### *Xenopus laevis n-cadherin* partial promoter identification

To identify the basic promoter of *Xenopus* n-cad we used the following strategy. The basic promoter region from chick and mammalian *n-cadherin* genes have been described at the 5’-UTR regions, within the position -3000 to -1 bp respect the translation initiation site (Li et al 1997; Mee et al 2005). By using the *Xenopus laevis* Genome Project resource (https://xenopus.lab.nig.ac.jp/) we identified a region of 2800 bp in the 5’-UTR of the *Xenopus laevis n-cadherin* gene (Figure S7). Since our data indicate that Cx43Tail, BTF-3, and Pol II form a complex (Figure 7C), we use ElemeNT tool (Sloutskin et al., 2015) to search for potentially active TATA boxes in the region that we isolated. Our *in silico* analysis reveal a TATA box rich region between the positions -166 to -618 bp, relative to the translation start site (Figure S7). Finally, we design overlapping primers to amplify fragments of 200 bp across this region (Figure S7). Primers sequence are listed in the ChIP section and their binding regions are hihglited in Figure S7.

### Chromatin immunoprecipitation (ChIP)

For chromatin immunoprecipitation (Chip), we followed a standard procedure for Xenopus laevis embryos (Gentsch and Smith, 2014), using 15 min for fixation of neurula stages Xenopus laevis embryos and 3 μg of Pol II antibody (Diagenode, C15100055) or GFP ChIP grade antibody (Abcam, ab290). For DNA extraction we followed a standard protocol (Akkers et al., 2012). Using the ElemeNT analysis resource we examined the Xenopus laevis n-cad promoter region for putative TATA boxes (FigureS7A) and designed primers to analyse the Chip samples by PCR. PCR was performed using the following protocol 95°C for 30 s, 56°C for 40 s and 72°C for 30 s for 32 cycles. Primers used for the Chip-PCR are: P5F: 5’-CTTCCAAGAGATGAAGCTCATAT-3’, P5R: 5’- AACACTCTATATGGCAGATAAC-3’, P6F: 5’-CCTTTAAATGCATACACTTACC-3’, P6R: 5’-

ACAGAAAAAGCATTTGCTTCCT-3’, P7F: 5’-CAATCAGATCCTTATATGTCCC-3’, P7R: 5’-GCCAAGTTTTCCCTTTGTTGT-3’, P8F: 5’-

GGAAGCAAATGCTTTTTCTGTC-3’, P8R: 5’-AGTCTGCTTTAGGAGACAACG-3’ and their relative binding sites are shown in FigureS7B.

## Supplemantal Movies

**Movie S1. Cx43 is required for collective NC migration. Related to Figure1.**

Example of chemotaxis assay with CTLMO NC explants (left side) displaying directional migration towards the chemotactic cue SDF-1 and Cx43MO NC explants that do not migrate towards the SDF-1 bead (right side). Green is membrane-GFP and red/orange is nuclear-mCherry. Frame delay 10 min. Magnification 10X.

**Movie S2. Cx43 is not involved in NC cell motility. Related to Figure1.**

Example of single cell tracks with CTLMO NC cells in cyan (left side) and Cx43MO in yellow(right side) and individual cell teacks. Cyan and yellow are membrane-GFP and magenta is nuclear-mCherry. Frame delay 5 min. Magnification 10X.

**Movie S3. Cx43 depletion inhibits CIL. Related to Figure2.**

Example of cell collision assay with CTLMO NC cells undergoing CIL (left side) and Cx43MO NC cells that stay longer in contact and do not exhibit CIL (right side). Cyan and yellow are membrane-GFP and magenta is nuclear-mCherry. Arrows display direction of each cell that is tracked. Frame delay 5 min. Magnification 20X.

**Movie S4. Cx43 inhibition delays NC explant dispersion in vitro. Related to Figure2.**

Example of cell dispersion assay with CTLMO NC explants dispersing over time(left side) and Cx43MO NC explants showing a later dispersion (right side). Red is nuclear-mCherry. Frame delay is 10 min. Magnfication 20X.

**Movie S5. Perturbation of cell dispersion in Cx43MO explants is rescued by n-cad. Related to Figure3.**

Example of cell dispersion assay. After 6hr of culture, CTLMO NC explants disperse (left), Cx43MO NC explants do not disperse (middle) and Cx43MO+n-cad mRNA NC explants disperse (right). Red is nuclear-mCherry. Frame delay is 1 hr. Magnfication 20X.

## References

Ai, Z., Fischer, A., Spray, D.C., Brown, A.M.C., and Fishman, G.I. (2000). Wnt-1 regulation of connexin43 in cardiac myocytes. J. Clin. Invest. 105, 161–171.

Akkers, R.C., Jacobi, U.G., and Veenstra, G.J.C. (2012). Chromatin immunoprecipitation analysis of Xenopus embryos. Methods Mol. Biol. 917, 279–292.

Alexander, D.B., and Goldberg, G.S. (2003). Transfer of biologically important molecules between cells through gap junction channels. Curr. Med. Chem. 10, 2045–2058.

Alfandari, D., Cousin, H., Gaultier, A., Hoffstrom, B.G., and DeSimone, D.W. (2003). Integrin α5β1 supports the migration of Xenopus cranial neural crest on fibronectin. Dev. Biol. 260, 449–464.

Aybar, M.J., Nieto, M.A., and Mayor, R. (2003). Snail precedes slug in the genetic cascade required for the specification and migration of the Xenopus neural crest. Development 130, 483–494.

Barriga, E.H., Maxwell, P.H., Reyes, A.E., and Mayor, R. (2013). The hypoxia factor Hif-1?? controls neural crest chemotaxis and epithelial to mesenchymal transition. J. Cell Biol. 201, 759–776.

Borchers, a, Epperlein, H.H., and Wedlich, D. (2000). An assay system to study migratory behavior of cranial neural crest cells in Xenopus. Dev. Genes Evol. 210, 217–222.

Bruzzone, S., Guida, L., Zocchi, E., Franco, L., and De Flora A (2001). Connexin 43 hemi channels mediate Ca2+-regulated transmembrane NAD+ fluxes in intact cells. FASEB J. 15, 10–12.

De Calisto, J., Araya, C., Marchant, L., Riaz, C.F., and Mayor, R. (2005). Essential role of non-canonical Wnt signalling in neural crest migration. Development 132, 2587–2597.

Carmona-Fontaine, C., Matthews, H.K., Kuriyama, S., Moreno, M., Dunn, G. a., Parsons, M., Stern, C.D., and Mayor, R. (2008). Contact inhibition of locomotion in vivo controls neural crest directional migration. Nature 456, 957–961.

Cavallini, B., Huet, J., Plassat, J.L., Sentenac, A., Egly, J.M., and Chambon, P. (1988). A yeast activity can substitute for the HeLa cell TATA box factor. Nature 334, 77–80.

Chakrabarti, a, Matthews, G., Colman, a, and Dale, L. (1992). Secretory and inductive properties of Drosophila wingless protein in Xenopus oocytes and embryos. Dev. Cambridge Engl. 115, 355–369.

Cina, C., Maass, K., Theis, M., Willecke, K., Bechberger, J.F., and Naus, C.C. (2009). Involvement of the cytoplasmic C-terminal domain of connexin43 in neuronal migration. J. Neurosci. 29, 2009–2021.

Dent, J.A., Polson, A.G., and Klymkowsky, M.W. (1989). A whole-mount immunocytochemical analysis of the expression of the intermediate filament protein vimentin in Xenopus. 74, 61–74.

Dimauro, I., Pearson, T., Caporossi, D., and Jackson, M.J. (2012). A simple protocol for the subcellular fractionation of skeletal muscle cells and tissue. BMC Res. Notes 5, 513.

Ebihara, L., Xu, X., Oberti, C., Beyer, E.C., and Berthoud, V.M. (1999). Co-Expression of Lens Fiber Connexins Modifies Hemi-Gap-Junctional Channel Behavior. 76, 198–206.

Elias, L.A.B., Wang, D.D., and Kriegstein, A.R. (2007). Gap junction adhesion is necessary for radial migration in the neocortex. Nature 448, 901–907.

Elias, L.A.B., Turmaine, M., Parnavelas, J.G., and Kriegstein, A.R. (2010). Connexin 43 mediates the tangential to radial migratory switch in ventrally derived cortical interneurons. J. Neurosci. 30, 7072–7077.

Elzarrad, M.K., Haroon, A., Willecke, K., Dobrowolski, R., Gillespie, M.N., and Al-Mehdi, A.-B. (2008). Connexin-43 upregulation in micrometastases and tumor vasculature and its role in tumor cell attachment to pulmonary endothelium. BMC Med. 6, 20.

Ewart, J.L., Cohen, M.F., Meyer, R.A., Huang, G.Y., Wessels, A., Gourdie, R.G., and Chin, A.J. (1997). Heart and neural tube defects in transgenic mice overexpressing the Cx43 gap junction gene. 1292, 1281–1292.

Fawcett, S.R., and Klymkowsky, M.W. (2004). Embryonic expression of Xenopus laevis SOX7. Gene Expr. Patterns 4, 29–33.

Francis, R., Xu, X., Park, H., Wei, C.J., Chang, S., Chatterjee, B., and Lo, C. (2011). Connexin43 modulates cell polarity and directional cell migration by regulating microtubule dynamics. PLoS One 6, e26379. doi: 10.1371/journal.pone.0026379.

Friedl, P., and Gilmour, D. (2009). Collective cell migration in morphogenesis, regeneration and cancer. Nat. Rev. Mol. Cell Biol. 10, 445–457.

Fu, C.T., Bechberger, J.F., Ozog, M.A., Perbal, B., and Naus, C.C. (2004). CCN3 (NOV) interacts with connexin43 in C6 glioma cells: Possible mechanism of connexin-mediated growth suppression. J. Biol. Chem. 279, 36943–36950.

Gentsch, G.E., and Smith, J.C. (2014). Investigating physical chromatin associations across the xenopus genome by chromatin immunoprecipitation. Cold Spring Harb. Protoc. 2014, 483–497.

Gentzel, M., Schille, C., Rauschenberger, V., and Schambony, A. (2015). Distinct functionality of dishevelled isoforms on Ca2+/calmodulin-dependent protein kinase 2 (CamKII) in Xenopus gastrulation. Mol. Biol. Cell 26, 966–977.

Giepmans, B.N.G. (2006). Role of Connexin43-Interacting Proteins at Gap Junctions. 42, 41–56.

Giepmans, B.N.G., Verlaan, I., Hengeveld, T., Janssen, H., Calafat, J., Falk, M.M., and Moolenaar, W.H. (2001). Gap junction protein connexin-43 interacts directly with microtubules. Curr. Biol. 11, 1364–1368.

Gupta, N., Wang, H., McLeod, T.L., Naus, C.C., Kyurkchiev, S., Advani, S., Yu, J., Perbal, B., and Weichselbaum, R.R. (2001). Inhibition of glioma cell growth and tumorigenic potential by CCN3 (NOV). Mol. Pathol. 54, 293–299.

Harland, R. (1991). In situ hybridization: an improved whole-mount method for Xenopus embryos. Methods Cell Biol. 36.

Harland, R., and Weintraub, H. (1985). Translation of mRNA injected into Xenopus oocytes is specifically inhibited by antisense RNA. J. Cell Biol. 101, 1094–1099.

Homkajorn, B., Sims, N.R., and Muyderman, H. (2010). Connexin 43 regulates astrocytic migration and proliferation in response to injury. Neurosci. Lett. 486, 197–201.

Hong, C.-S., and Saint-Jeannet, J.-P. (2007). The activity of Pax3 and Zic1 regulates three distinct cell fates at the neural plate border. Mol. Biol. Cell 18, 2192–2202.

Hopwood, N.D., Pluck, A., and Gurdon, J.B. (1989). A Xenopus mRNA related to Drosophila twist is expressed in response to induction in the mesoderm and the neural crest. Cell 59, 893–903.

Kang, E.Y., Ponzio, M., Gupta, P.P., Liu, F., Butensky, A., and Gutstein, D.E. (2009). Identification of binding partners for the cytoplasmic loop of connexin43: a novel interaction with β-tubulin. Cell Commun. Adhes. 15, 397–406.

Kashef, J., Köhler, A., Kuriyama, S., Alfandari, D., Mayor, R., and Wedlich, D. (2009). Cadherin-11 regulates protrusive activity in Xenopus cranial neural crest cells upstream of Trio and the small GTPases. Genes Dev. 23, 1393–1398.

Krause, M., Leslie, J.D., Stewart, M., Lafuente, E.M., Valderrama, F., Jagannathan, R., Strasser, G. a, Rubinson, D. a, Liu, H., Way, M., et al. (2004). Lamellipodin, an Ena/VASP ligand, is implicated in the regulation of lamellipodial dynamics. Dev. Cell 7, 571–583.

Kushima, S., Mammadova, G., Mahbub Hasan, a K.M., Fukami, Y., and Sato, K. (2011). Characterization of lipovitellin 2 as a tyrosine-phosphorylated protein in oocytes, eggs and early embryos of Xenopus laevis. Zoolog. Sci. 28, 550–559.

Kusumawidjaja, G., Kayed, H., Giese, N., Bauer, A., Erkan, M., Giese, T., Hoheisel, J.D., Friess, H., and Kleeff, J. (2007). Basic transcription factor 3 (BTF3) regulates transcription of tumor-associated genes in pancreatic cancer cells. Cancer Biol. Ther. 6, 367–376.

Kwak, B.R., Hermans, M.M., De Jonge, H.R., Lohmann, S.M., Jongsma, H.J., and Chanson, M. (1995). Differential regulation of distinct types of gap junction channels by similar phosphorylating conditions. Mol. Biol. Cell 6, 1707–1719.

Law, A.L., Vehlow, A., Kotini, M., Dodgson, L., Soong, D., Theveneau, E., Bodo, C., Taylor, E., Navarro, C., Perera, U., et al. (2013). Lamellipodin and the Scar/WAVE complex cooperate to promote cell migration in vivo. J. Cell Biol. 203, 673–689.

Linker, C., Bronner-Fraser, M., and Mayor, R. (2000). Relationship between gene expression domains of Xsnail, Xslug, and Xtwist and cell movement in the prospective neural crest of Xenopus. Dev. Biol. 224, 215–225.

Liu, X., Sun, L., Torii, M., and Rakic, P. (2012). Connexin 43 controls the multipolar phase of neuronal migration to the cerebral cortex. Proc. Natl. Acad. Sci. U. S. A. 109, 8280–8285.

Mathias, R.T., White, T.W., and Gong, X. (2010). Lens gap junctions in growth, differentiation, and homeostasis. Physiol. Rev. 90, 179–206.

Matthews, H.K., Marchant, L., Carmona-Fontaine, C., Kuriyama, S., Larraín, J., Holt, M.R., Parsons, M., and Mayor, R. (2008). Directional migration of neural crest cells in vivo is regulated by Syndecan-4/Rac1 and non-canonical Wnt signaling/RhoA. Development 135, 1771–1780.

Mayor, R., Morgan, R., and Sargent, M.G. (1995). Induction of the prospective neural crest of Xenopus. 777, 767–777.

Mendoza-Naranjo, A., Cormie, P., Serrano, A.E., Wang, C.M., Thrasivoulou, C., Sutcliffe, J.E.S., Gilmartin, D.J., Tsui, J., Serena, T.E., Phillips, A.R.J., et al. (2012). Overexpression of the gap junction protein Cx43 as found in diabetic foot ulcers can retard fibroblast migration. Cell Biol. Int. 36, 661–667.

Moore, R., Theveneau, E., Pozzi, S., Alexandre, P., Richardson, J., Merks, A., Parsons, M., Kashef, J., Linker, C., and Mayor, R. (2013). Par3 controls neural crest migration by promoting microtubule catastrophe during contact inhibition of locomotion. Development 140, 4763–4775.

Nandadasa, S., Tao, Q., Menon, N.R., Heasman, J., and Wylie, C. (2009). N- and E-cadherins in Xenopus are specifically required in the neural and non-neural ectoderm, respectively, for F-actin assembly and morphogenetic movements. Development 136, 1327–1338.

Newport, J., and Kirschner, M. (1982). A major developmental transition in early xenopus embryos: I. characterization and timing of cellular changes at the midblastula stage. Cell 30, 675–686.

Niessen, H., Harz, H., Bedner, P., Krämer, K., Willecke, K., Genetik, I., Molekulargenetik, A., Bonn, U., Institut, B., and Straße, M. (2006). Selective permeability of different connexin channels to the second messenger cyclic AMP. J. Biol. Chem. 281, 6673–6681.

Nieuwkoop P., F.J. (1994). Normal Table of Xenopus Laevis (Daudin): A Systematical & Chronological Survey of the Development from the Fertilized Egg till the End of Metamorphosis.

Nones, J., Costa, A.P., Leal, R.B., Gomes, F.C.A., and Trentin, A.G. (2012). The flavonoids hesperidin and rutin promote neural crest cell survival. Cell Tissue Res. 350, 305–315.

Ogawa, K., Pitchakarn, P., Suzuki, S., Chewonarin, T., Tang, M., Takahashi, S., Naiki-Ito, A., Sato, S., Takahashi, S., Asamoto, M., et al. (2012). Silencing of connexin 43 suppresses invasion, migration and lung metastasis of rat hepatocellular carcinoma cells. Cancer Sci. 103, 860–867.

Plum, A., Hallas, G., Magin, T., Dombrowski, F., Hagendorff, A., Schumacher, B., Wolpert, C., Kim, J.S., Lamers, W.H., Evert, M., et al. (2000). Unique and shared functions of different connexins in mice. Curr. Biol. 10, 1083–1091.

Saidi Brikci-Nigassa, A., Clement, M.J., Ha-Duong, T., Adjadj, E., Ziani, L., Pastre, D., Curmi, P.A., and Savarin, P. (2012). Phosphorylation controls the interaction of the connexin43 C-terminal domain with tubulin and microtubules. Biochemistry 51, 4331–4342.

Saka, Y., Hagemann, A.I., Piepenburg, O., and Smith, J.C. (2007). Nuclear accumulation of Smad complexes occurs only after the midblastula transition in Xenopus. Development 134, 4209–4218.

Salat-Canela, C., Sesé, M., Peula, C., Ramón, y Cajal, S., and Aasen, T. (2014). Internal translation of the connexin 43 transcript. Cell Commun. Signal. 12, 31.

Scarpa, E., Szabó, A., Bibonne, A., Theveneau, E., Parsons, M., and Mayor, R. (2015). Cadherin Switch during EMT in Neural Crest Cells Leads to Contact Inhibition of Locomotion via Repolarization of Forces. Dev. Cell 421–434.

Smyth, J.W., and Shaw, R.M. (2013). Autoregulation of connexin43 gap junction formation by internally translated isoforms. Cell Rep. 5, 611–618.

Solan, J.L., and Lampe, P.D. (2005). Connexin phosphorylation as a regulatory event linked to gap junction channel assembly. Biochim. Biophys. Acta 1711, 154–163.

Steventon, B., Araya, C., Linker, C., Kuriyama, S., and Mayor, R. (2009). Differential requirements of BMP and Wnt signalling during gastrulation and neurulation define two steps in neural crest induction. Development 136, 771–779.

Stoletov, K., Strnadel, J., Zardouzian, E., Momiyama, M., Park, F.D., Kelber, J.A., Pizzo, D.P., Hoffman, R., VandenBerg, S.R., and Klemke, R.L. (2013). Role of connexins in metastatic breast cancer and melanoma brain colonization. J. Cell Sci. 126, 904–913.

Stramer, B.M., Dunn, G.A., Davis, J.R., and Mayor, R. (2013). Rediscovering contact inhibition in the embryo. J. Microsc. 251, 206–211.

Taillard, É.D., Waelti, P., and Zuber, J. (2008). Few statistical tests for proportions comparison. Eur. J. Oper. Res. 185, 1336–1350.

Takamatsu, T., Regulation, C., and Prefectural, K. (2007). Cx43 Mediates TGF-_NL_ Signaling through Competitive Smads Binding to Microtubules ▭. 18, 2264–2273.

Theveneau, E., and Mayor, R. (2011). Beads on the run: Beads as alternative tools for chemotaxis assays. Methods Mol. Biol. 769, 449–460.

Theveneau, E., and Mayor, R. (2013). Collective cell migration of epithelial and mesenchymal cells. Cell. Mol. Life Sci. 70, 3481–3492.

Theveneau, E., Marchant, L., Kuriyama, S., Gull, M., Moepps, B., Parsons, M., and Mayor, R. (2010). Collective chemotaxis requires contact-dependent cell polarity. Dev. Cell 19, 39–53.

Theveneau, E., Steventon, B., Scarpa, E., Garcia, S., Trepat, X., Streit, A., and Mayor, R. (2013). Chase-and-run between adjacent cell populations promotes directional collective migration. Nat. Cell Biol. 15, 763–772.

Tribulo, C., Aybar, M.J., Nguyen, V.H., Mullins, M.C., and Mayor, R. (2003). Regulation of Msx genes by a Bmp gradient is essential for neural crest specification. Development 130, 6441–6452.

Ul-Hussain, M., Olk, S., Schoenebeck, B., Wasielewski, B., Meier, C., Prochnow, N., May, C., Galozzi, S., Marcus, K., Zoidl, G., et al. (2014). Internal ribosomal entry site (IRES) activity generates endogenous carboxyl-terminal domains of Cx43 and is responsive to hypoxic conditions. J. Biol. Chem. 289, 20979–20990.

Waldo, K.L., Lo, C.W., and Kirby, M.L. (1999). Connexin 43 expression reflects neural crest patterns during cardiovascular development. Dev. Biol. 208, 307–323.

Wang, W., Xu, M., Wang, Y., and Jamil, M. (2014). Basal transcription factor 3 plays an important role in seed germination and seedling growth of rice. Biomed Res. Int. 2014.

Xu, X. (2001). Modulation of mouse neural crest cell motility by N-cadherin and connexin 43 gap junctions. J. Cell Biol. 154, 217–230.

Xu, X., Li, W.E., Huang, G.Y., Meyer, R., Chen, T., Luo, Y., Thomas, M.P., Radice, G.L., and Lo, C.W. (2001). Modulation of mouse neural crest cell motility by N-cadherin and connexin 43 gap junctions. J. Cell Biol. 154, 217–230.

Xu, X., Francis, R., Wei, C.J., Linask, K.L., and Lo, C.W. (2006). Connexin 43-mediated modulation of polarized cell movement and the directional migration of cardiac neural crest cells. Development 133, 3629–3639.

Zhang, X., Sun, Y., Wang, Z., Huang, Z., Li, B., and Fu, J. (2014). Up-regulation of connexin-43 expression in bone marrow mesenchymal stem cells plays a crucial role in adhesion and migration of multiple myeloma cells. Leuk. Lymphoma 1–8.

Zheng, X.M., Moncollin, V., Egly, J.M., and Chambon, P. (1987). A general transcription factor forms a stable complex with RNA polymerase B (II). Cell 50, 361–368.

Zheng, X.M., Black, D., Chambon, P., and Egly, J.M. (1990). Sequencing and expression of complementary DNA for the general transcription factor BTF3. Nature 344.

